# A paracrine circuit of IL-1β/IL-1R1 between myeloid and tumor cells drives glioblastoma progression

**DOI:** 10.1101/2022.04.03.486888

**Authors:** Zhihong Chen, Bruno Giotti, Milota Kaluzova, Cameron J. Herting, Gonzalo Pinero, Montse Puigdelloses Vallcorba, Simona Cristea, James L. Ross, James Ackley, Victor Maximov, Frank Szulzewsky, Mar Marquez-Ropero, Angelo Angione, Noah Nichols, Nadejda Tsankova, Franziska Michor, Dmitry M. Shayakhmetov, David H. Gutmann, Alexander M. Tsankov, Dolores Hambardzumyan

## Abstract

Monocytes and monocyte-derived macrophages (MDM) from blood circulation infiltrate and promote glioblastoma growth. Here we discover that glioma cells induce the expression of potent pro-inflammatory cytokine IL-1β in MDM, which engages IL-1R1 in glioma cells, activates NF-κB pathway, and subsequently leads to the induction of monocyte chemoattractant proteins (MCPs). Thus, a feedforward paracrine circuit of IL-1β/IL-1R1 between the tumors and MDM creates an interdependence driving glioblastoma progression. Locally antagonizing IL-1β/IL-1R1 leads to reduced MDM infiltration, diminished tumor growth, reduced exhausted CD8^+^ T cells, and thereby extends the survival of tumor-bearing mice. In contrast to IL-1β, IL-1a exhibits anti-tumor effects. Genetic deletion of *Il1a* is associated with decreased recruitment of lymphoid cells and loss of interferon (IFN) signaling in various immune populations and subsets of malignant cells. IL-1β antagonism of IL-1β should be considered as an effective anti-glioblastoma therapy.

**Graphical Abstract:** 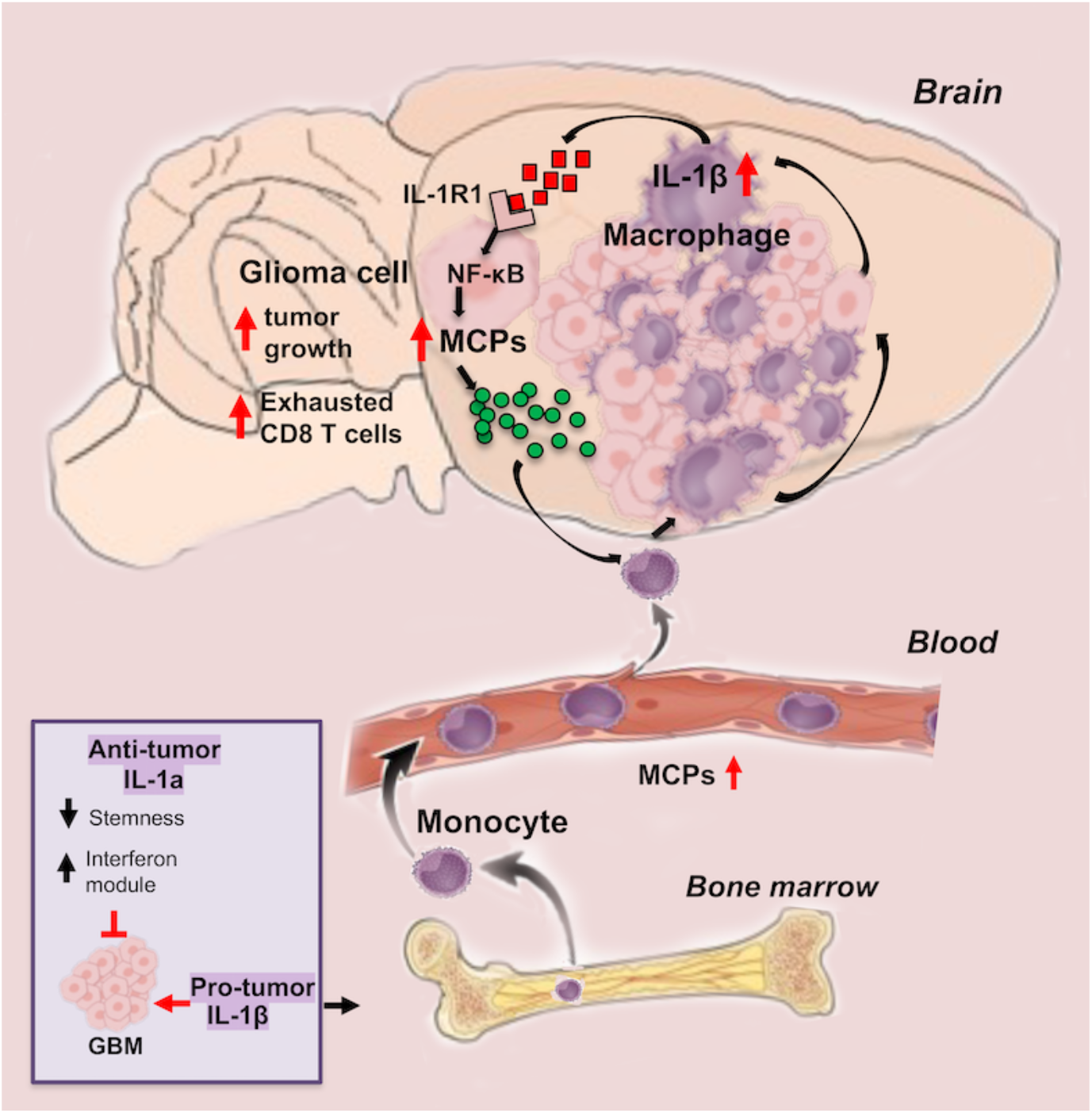

## Introduction

Glioblastoma (GBM) is the most prevalent and aggressive primary brain tumor in adults, associated with overall poor survival rates. Over the years, numerous therapies have entered clinical trials, with only temozolomide (TMZ) and radiotherapy (RT) modestly increasing median survival (Stupp et al., 2005). One reason that current anti-neoplastic therapies have not significantly improved patient outcomes is that these tumors are composed of numerous non-neoplastic cells that drive cancer progression. These non-neoplastic cells in the tumor microenvironment (TME) have the capacity to change as a function of tumor evolution and treatment, creating highly adaptive cancers with the ability to evade tumor-directed therapies. The non-neoplastic cells in the GBM TME include myeloid cells of innate immune system, with a predominance of tumor-associated macrophages (TAMs), which originate either from the bone marrow (bone marrow-derived myeloid cells, BMDM) or resident brain intrinsic microglia (MG) (Bowman et al., 2016; Chen et al., 2017). Within this myeloid compartment, there is a high degree of intra- and inter-tumor heterogeneity, representing regional differences in composition and function within tumors, as well as non-mutually exclusive subclass differences between mesenchymal (MES), proneural (PN), and classical (CL) GBM (Kaffes et al., 2019; Wang et al., 2018), that were determined based on dominant expression pattern at the time and location of tumor resection. Moreover, single-cell RNA sequencing analysis revealed that reciprocal interactions between TAMs and the tumor cells can drive a transition of GBM into a mesenchymal-like state (Hara et al., 2021). Thus, resistance of heterogeneous GBM with multiple cellular states to current standard-of-care therapy may be improved with TAM-targeted immunotherapies. (Chen and Hambardzumyan, 2021; Hara et al., 2021).

While TAMs are critical for glioma progression (Chatterjee et al., 2021), they are generally regarded as “immunosuppressive”, characterized by the release of a wide array of growth factors, chemokines, and cytokines in response to factors produced by tumor cells (Buonfiglioli and Hambardzumyan, 2021; Hambardzumyan et al., 2016). One such cytokine produced by BMDM is the master pro-inflammatory regulator, interleukin-1 (IL-1 denotes both IL-1α and IL-1β), which induces GBM-associated vasogenic edema (Herting et al., 2019). Increased expression of IL-1α and IL-1β has been reported in numerous cancers, where their tumor-promoting roles have also been established (Litmanovich et al., 2018). Although IL-1α and IL-1β both signal through the same ubiquitously-expressed receptor (IL-1R1), the resulting effects on tumor biology can be distinct and tissuespecific (Baker et al., 2019; Carmi et al., 2013; Fahey and Doyle, 2019).

To determine whether IL-1α and IL-1β have differential effects on GBM development, we leveraged the *RCAS/Ntv-a* somatic cell type-specific gene transfer system to create a series of genetically-engineered mouse models (GEMM), imitating the molecular subtypes of human GBM (Herting et al., 2017). Using these murine GBM (mGBM) models, we investigated how IL-1α and IL-1β contribute to tumor development *in vitro* and *in vivo*. We describe a paracrine circuit in which GBM cells recruit and induce IL-1 expression in BMDMs, which results in NF-κB pathway activation in GBM cells, leading to increased expression of the monocyte chemoattractant protein (MCP) and increased chemotaxis of inflammatory monocytes. In addition, we demonstrate that genetic loss of IL-1β leads to decreased inflammatory monocyte influx, reduced MG proliferation, and prolonged survival of mice. Similarly, pharmacologically inhibition of IL-1β or IL-1R1 decreases intratumoral TAM content and prolongs the survival of tumor-bearing mice. However, genetic deletion of both IL-1α and IL-1β results in decreased recruitment of lymphoid cells and loss of interferon (IFN) signaling in immune cells, allowing the tumor to escape immune control and thereby, worsening survival. Collectively, these findings establish a feedforward paracrine circuit of IL-1β/IL-1R1 between the BMDMs and tumors that drives glioblastoma progression. Local antagonism of IL-1β can be considered as an effective therapy for GBM.

## Results

### Increased IL1β expression in IDH1 WT GBM is associated with reduced patient survival

GBM tumors can be divided into two clinically distinct groups according to isocitrate dehydrogenase I and II (IDH 1/2) gene mutation status, where patients with mutant IDH (IDH-Mut) have longer survival times than those with wild type (WT) IDH (IDH-WT) (Louis et al., 2021; Parsons et al., 2008). We first evaluated *IL1A* and *IL1B* expression in IDH-WT and IDH-Mut samples using available TCGA datasets (U133 microarray data as detailed in the Method). While there were no differences in *IL1A* expression, *IL1B* expression was higher in IDH-WT samples (**Fig. 1A**, P<0.05), consistent with a recent report showing that IL-1β increases the growth of IDH-WT tumors (Liu et al., 2021). To determine whether IL-1 and its signaling pathway impact survival of patient with IDH-WT GBM, we used a Cox Proportional Hazards regression model to assess survival as a function of the following independent covariates: (1) each of the genes associated with IL1 signaling, (2) the entire pathway (specifically, the average normalized expression of all genes of the pathway, and (3) a subset of selected genes (the average normalized expression of *CASP1*, *IL1A*, *IL1B*, *IL1R1*, *IL1RAP*, *MYD88*, *TRAF6*, *TOLLIP*, *JUN*, *NFKB1*, highlighted in **Fig. S1**).

**Figure 1.**
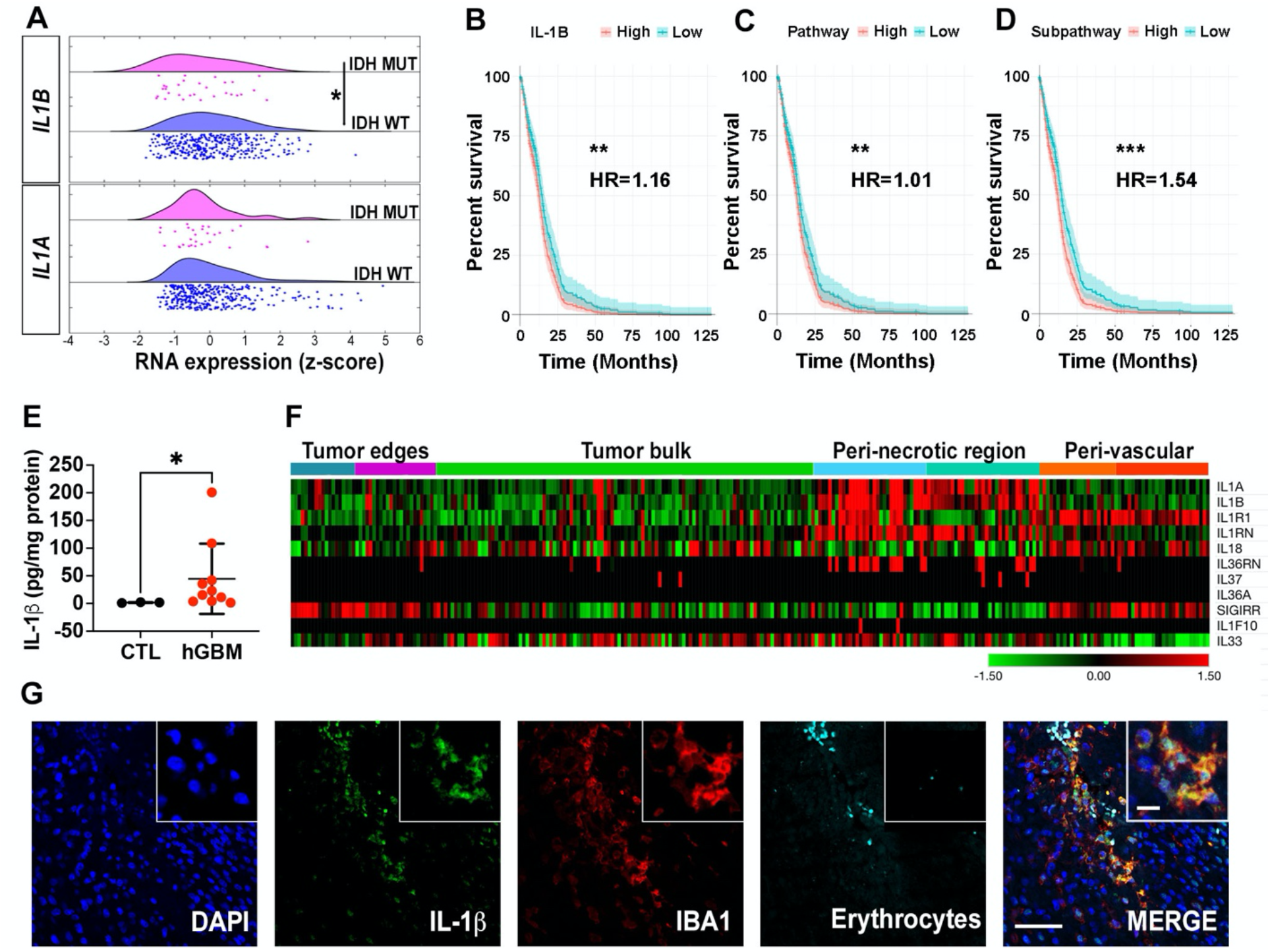
High *IL1B* expression in patients with IDH WT GBM is associated with reduced survival. (**A**) *IL1A* and *IL1B* expression was analyzed and compared in IDH WT (n=372) and IDH Mut (n=30) patient samples from TCGA datasets. Unpaired Student’s *t*-test, *P<0.05. (**B**) – (**D**) Survival curves grouped by high and low total expression (relative to the median) of the *IL1B* gene (**B**), IL-1 pathway (**C**) or a selected IL-1 subpathway (**D**, described in **SFig. 1**), fit using Cox Proportional Hazards regression models. Samples were culled from TCGA (n=372). P-values and HRs (Hazard ratio) come from the Cox Proportional Hazards Model including expression as a continuous covariate, and adjusting for gender and age. **P<0.01, ***P<0.001. (**E**) IL-1β ELISA in human IDH WT GBM samples. CTL: adjacent normal brain tissues. Unpaired Student’s *t*-test, *P<0.05. (**F**) Expression of *IL1* family members in various regions of human GBM tissues as defined by the IVYGap database (n=34) following laser capture microdissection and RNA-seq. (**G**) Representative images of immunofluorescence staining of IL-1 β (green), IBA1 (red), erythrocytes (cyan), and nuclei (visualized with DAPI, blue). Scale bar is 100 μm and 20 μm in insets.

The input data were culled from TCGA mRNA expression U133 microarray files. We included a total of 372 patient samples for which covariate information (survival information, age, and gender) was available. We found that both *IL-1B* gene (**Fig. 1B**, Hazard ratio (HR) >1, P<0.01) or IL1-related genes collectively serve as a significant adverse prognostic predictor of survival (**Fig. 1C**, HR >1, P<0.01). Particularly, when we analyzed the IL-1 signaling sub-pathway of selected genes that span the entire pathway, the HR increased to >1.5 (**Fig. 1D**, P=0.001), where four individual genes (*IL1A*, *IL1B*, *CASP1*, and *TOLLIP*) significantly influenced survival (**Fig. S1B-S1E**). We also found that the expression of the IL1 pathway and of any of its separate components does not impact survival differently depending on patient gender, as assessed by including an additional interaction term between gene expression and gender in the Cox Proportional Hazards Model (**Fig. S1B-S1E**). This contradicts a recent report showing that *IL1B* expression only correlated with patient survival when the patients were stratified by gender (Bayik et al., 2020). These discrepancies can be partially attributed to differences in cut-off values used to define high and low expressions, as well as the number of total samples utilized, and the statistical tests applied for the analyses.

Next, we confirmed increased IL-1β expression in microdissected IDH-WT human GBM tissues relative to adjacent normal brain tissues (**Fig. 1E**) (Sharma et al., 2011b). To determine where within the tumor *IL1B* and its receptor (*IL1RI*) are expressed, we queried the IVYGap (IVY Glioblastoma Atlas Project) database (https://glioblastoma.alleninstitute.org/), and found that *IL1B* and *IL1RI* are predominantly transcribed in peri-necrotic and peri-vascular regions, whereas the tumor bulk contains comparatively little expression (**Fig. 1F**). Additionally, we found that the majority of the *IL1* family members, including *IL1A* and *IL1Ra*, are similarly enriched in peri-necrotic regions (**Fig. 1F**), where we have previously shown that TAMs reside (Chen et al., 2017). By co-staining human GBM tissue sections for IL-1β and IBA1 (a pan-macrophage marker) expression, we found that IL-1β co-localizes with IBA1 in peri-vascular regions, suggesting that TAMs are a major source of IL-1β in human GBM (**Fig. 1G**).

### GBM cells induces IL-1 expression in BMDMs

Considering the co-localization of IL-1β and IBA1 in human GBM tissue, we next evaluated whether *IL1* expression levels differ between GBM subtypes. In light of previous studies demonstrating differences in TAM numbers and expression profiles between various human and murine GBM subtypes (Chen et al., 2020; Kaffes et al., 2019; Wang et al., 2018), we observed higher *IL1* expression levels in human MES, relative to PN, GBM (**Fig. 2A**, P<0.01). In addition, similar results were observed in *Nf1*-silenced murine GBM (model of human MES GBM, *Nf1* expression was suppressed by shRNA) compared to *PDGFB*-driven GBM (model of human PN GBM) (**Fig. 2B**, P<0.05). Consistent with the RNA expression results, IL-1β ELISA demonstrated that *Nf1*-silenced tumors had much higher IL-1β compared to *PDGFB*-driven tumors (**Fig. 2C**, P<0.01). This is not surprising since both murine *Nf1*-silenced GBM and human MES GBM exhibit significantly higher number of TAMs (Chen et al., 2020; Kaffes et al., 2019).

**Figure 2.**
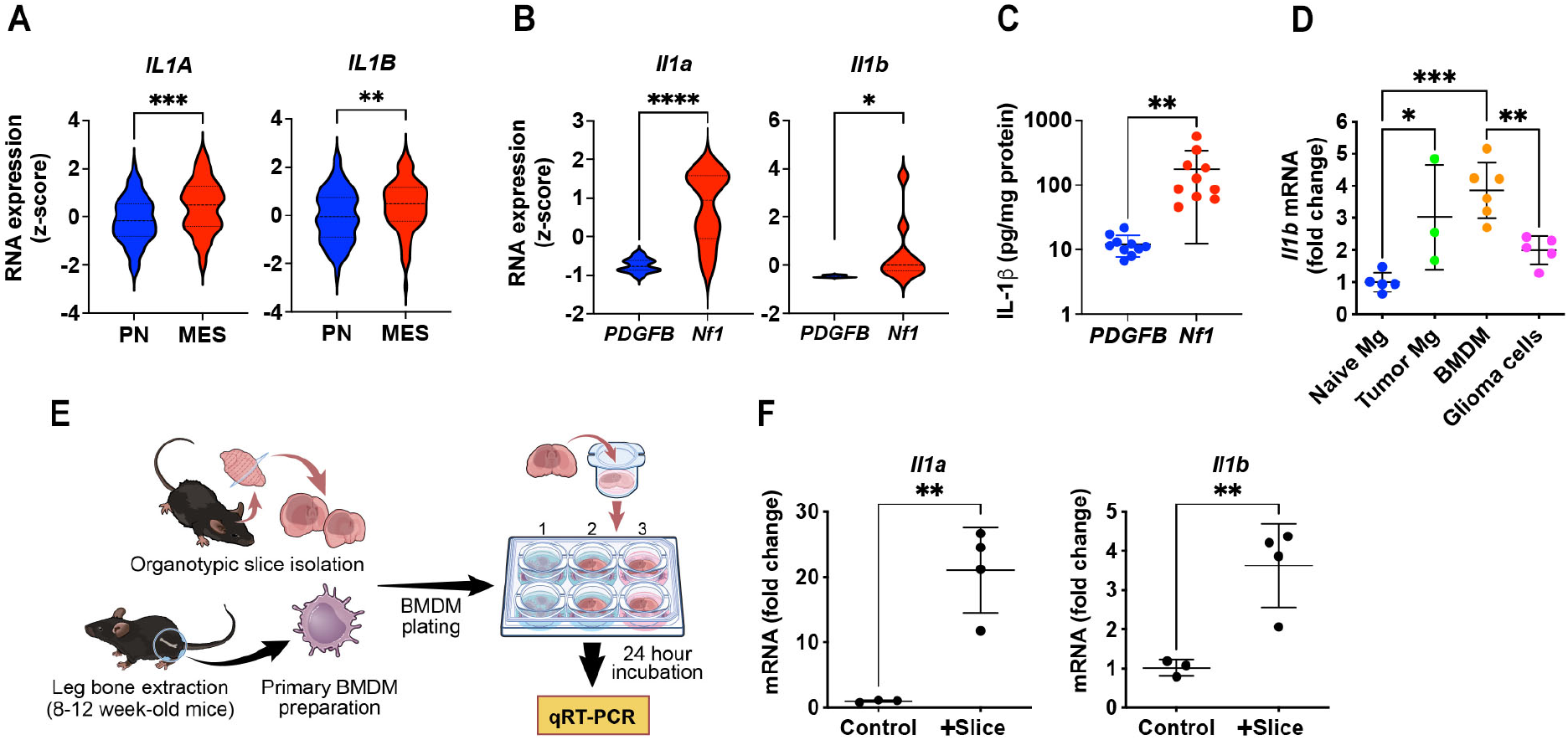
*IL1B* expression is increased in human MES GBM and *Nf1*-silenced murine GBM. (**A**) *IL1A* and *IL1B* RNA expression in PN and MES human GBM patient samples from TCGA. Two-tailed Student’s *t*-test, **P<0.01, ***P<0.001. (**B**) Quantitative PCR for *Il1a* and *Il1b* RNA expression from murine *PDGFB*-driven and *Nf1*-silenced GBM samples. Two-tailed Student’s *t*-test, *P<0.05, ****P<0.0001. (**C**) Quantification of IL-1β in *PDGFB*-driven and *Nf1*-silenced murine GBM tissues by ELISA. Two-tailed Student’s *t*-test, **P<0.01. (**D**) Quantitative PCR of *Il1b* expression in FACS-sorted cells from naïve brain and *PDGFB*-driven tumors. *P<0.05, **P<0.01, ***P<0.001, by one-way ANOVA with Tukey’s *post-hoc* comparisons. (**E**) Diagram illustrating the co-culturing system of primary murine BMDM and *PDGFB*-driven tumor slices. (**F**) Quantitative PCR of *Il1a* and *Il1b* expression in BMDM cocultured with tumor slices. **P<0.01, by two-tailed Student’s *t*-test.

To identify the source of IL-1β in these tumors, we used fluorescent-activated cell sorting (FACS) to isolate tumor-associated microglia, tumor-associated bone marrow derived macrophages (BMDM), and glioma cells for *Il1b* real-time quantitative PCR. Compared to naïve microglia from healthy control mice, all tumor-associated cell types showed increased *Il1b* mRNA (**Fig. 2D**), with BMDM demonstrating the highest levels of *Il1b* mRNA (**Fig. 2D**, P<0.001). To examine whether *Il1* expression in BMDM was induced by GBM tumor cells, naïve BMDMs were cocultured with primary murine *PDGFB*-driven GBM tumor slices (**Fig. 2E**). Both *Il1a* and *Il1b* were induced in BMDM by the tumor slices, but not by the medium used to maintain the slices (**Fig. 2F**). Together, these results establish that IL-1β is expressed in both human and murine GBMs, with greater levels in GBM of the MES subtype, and is likely induced in TAMs by GBM cells.

### *Il1b* genetic ablation prolongs tumor-bearing mouse survival

To determine whether IL-1β is essential for GBM growth, we generated *PDGFB*-driven primary tumors in wild type *(WT;Ntv-a)* and *Il1b-KO (Il1b^-/-^;Ntv-a)* mice (**Fig. 3A**) (Herting et al., 2017). In these KO mice, where *Il1b* was absent in both the tumor and TME cells, *Il1b* loss prolonged the survival of *PDGFB*-driven tumor-bearing mice (**Fig. 3A**). This prolonged survival was associated with decreased numbers of proliferating (phospho-histone 3-positive; pH3-positive) cells in the tumors (**Fig. 3B**).

**Figure 3.**
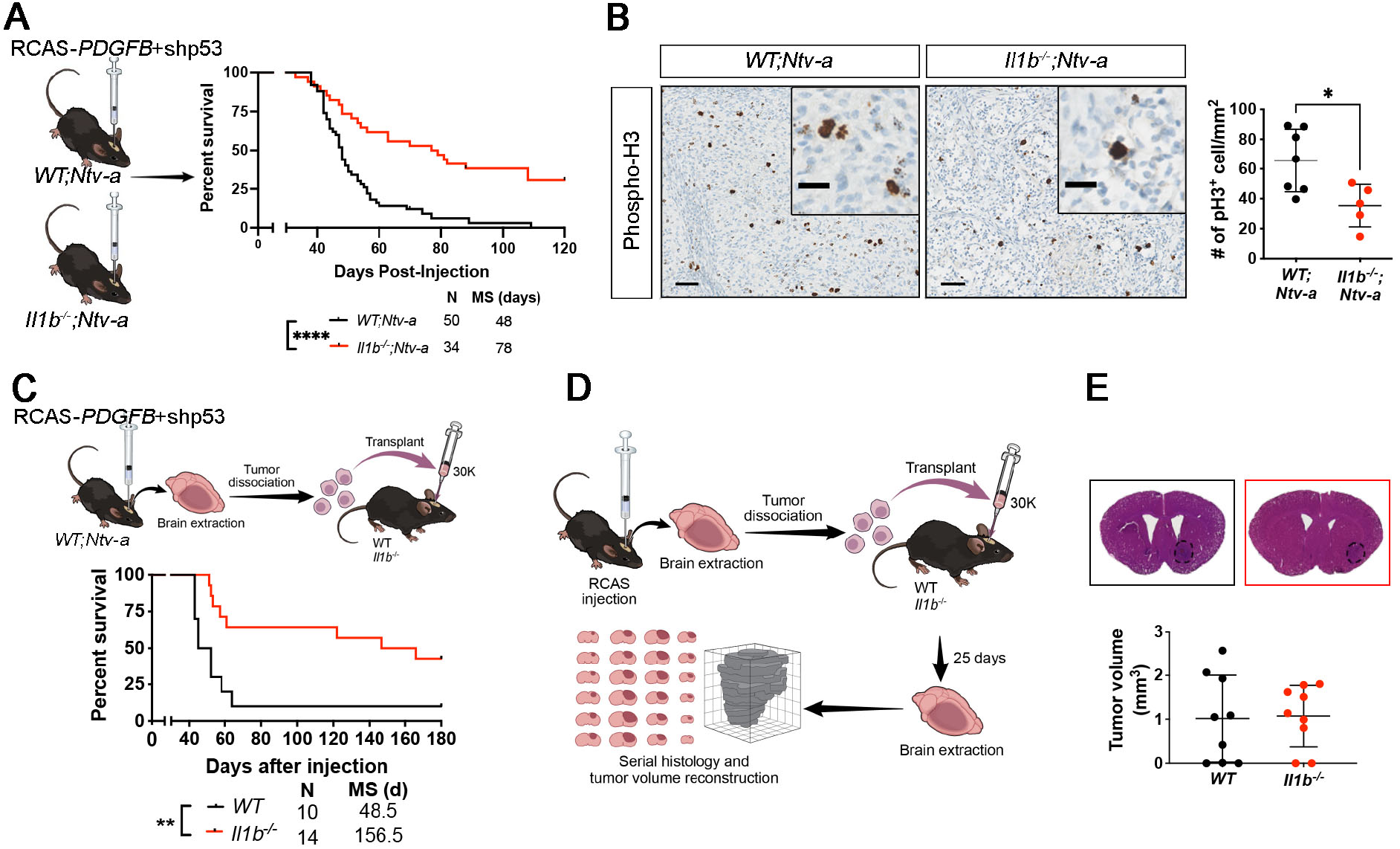
TME-derived IL-1β regulates *PDGFB-driven* murine GBM growth. (**A**) Schematic illustration of injections and Kaplan Meier-survival curves of *PDGFB*-driven tumors generated in *WT;Ntv-a* an *Il1b^-/-^;Ntv-a* mice. Curves were compared by log-rank (Mantel-Cox – MC) test, ****P<0.0001 and Gehan-Breslow-Wilcoxon (GBW) test, ***P<0.001 (not shown). (**B**) Representative pH3 immunohistochemistry images of *PDGFB*-driven tumors generated in *WT;Ntv-a* and *Il1b^-/-^;Ntv-a* mice. Two-tailed Student’s *t*-test, *P<0.05. Scale bar is 50 μm and 20 μm in insets. (**C**) Schematic illustration of orthotopic transplant of primary *PDGFB*-driven *Il1b* WT tumors into *Il1b^-/-^* and WT recipient animals, with the corresponding Kaplan-Meier survival curves. **P<0.01 by MC and GBW tests. (**D**) Schematic illustration of orthotopic transplant of primary *PDGFB*-driven *Il1* WT tumors into WT and *Il1b^-/-^* recipient animals to determine the effects on tumor volume during early tumor evolution. (**E**) Comparison of tumor volumes during early tumor evolution between tumors transplanted in WT and *Il1b^-/-^* recipient animals.

Next, to distinguish between prolonged survival reflecting *Il1b* loss in tumor versus TME cells, we *orthotopically* transplanted 30,000 cells from primary *PDGFB*-driven GBM (*WT;Ntv-a* mice) into the striatum of *WT* and *Il1b^-/-^* mice (**Fig. 3C**). Genetic *Il1b* ablation in the TME significantly prolonged the survival of tumor-bearing mice, recapitulating the results of the RCAS/*Ntv-a* system, and suggesting that this beneficial effect of *Il1b* loss is a TME-driven effect (**Fig. 3C**). To exclude defective tumor initiation in *Il1b* mice, mice were euthanized at an earlier time point (25 days post-tumor cell transplantation) and their brains serially sectioned for tumor volume estimates (Herting et al., 2019) (**Fig. 3D**). We found that the tumor volumes were comparable between *WT* and *Il1b* recipient mice at this early stage of tumor development, indicating that tumor initiation was not affected in *Il1b^-/-^* mice, but rather that reduced tumor growth in *Il1b^-/-^* mice is the driver of the prolonged survival (**Fig. 3E**).

### *Il1* loss decreases inflammatory monocyte tumor infiltration

To determine whether there is a functional redundancy between IL-1a and IL-1β, we analyzed tumors isolated from *WT;Ntv-a, Il1b^-/-^; Ntv-a*, and *Il1a^-/-^;Il1b^-/-^;Ntv-a* mice (**Fig. 4A**) by single-cell RNA sequencing (scRNA-seq) (**Fig. S2A**). Unsupervised clustering identified five major cell classes in the tumors: lymphoid and myeloid immune cells, stromal cells, endothelial cells, and tumor cells (**Fig. 4B**), with myeloid cells accounting for the majority of the non-malignant cell population, in agreement with our previous work (Chen et al., 2017). At a single cell level, myeloid cells were the major producers of *Il1a* and *Il1b*, while the *Rfp^+^* (red fluorescent protein, result of the RCAS-shp53-*Rfb*) transformed neoplastic cells do not transcribe these cytokines (**Fig. 4C**), in keeping with our immunostaining results (**Fig. 1E**).

**Figure 4.**
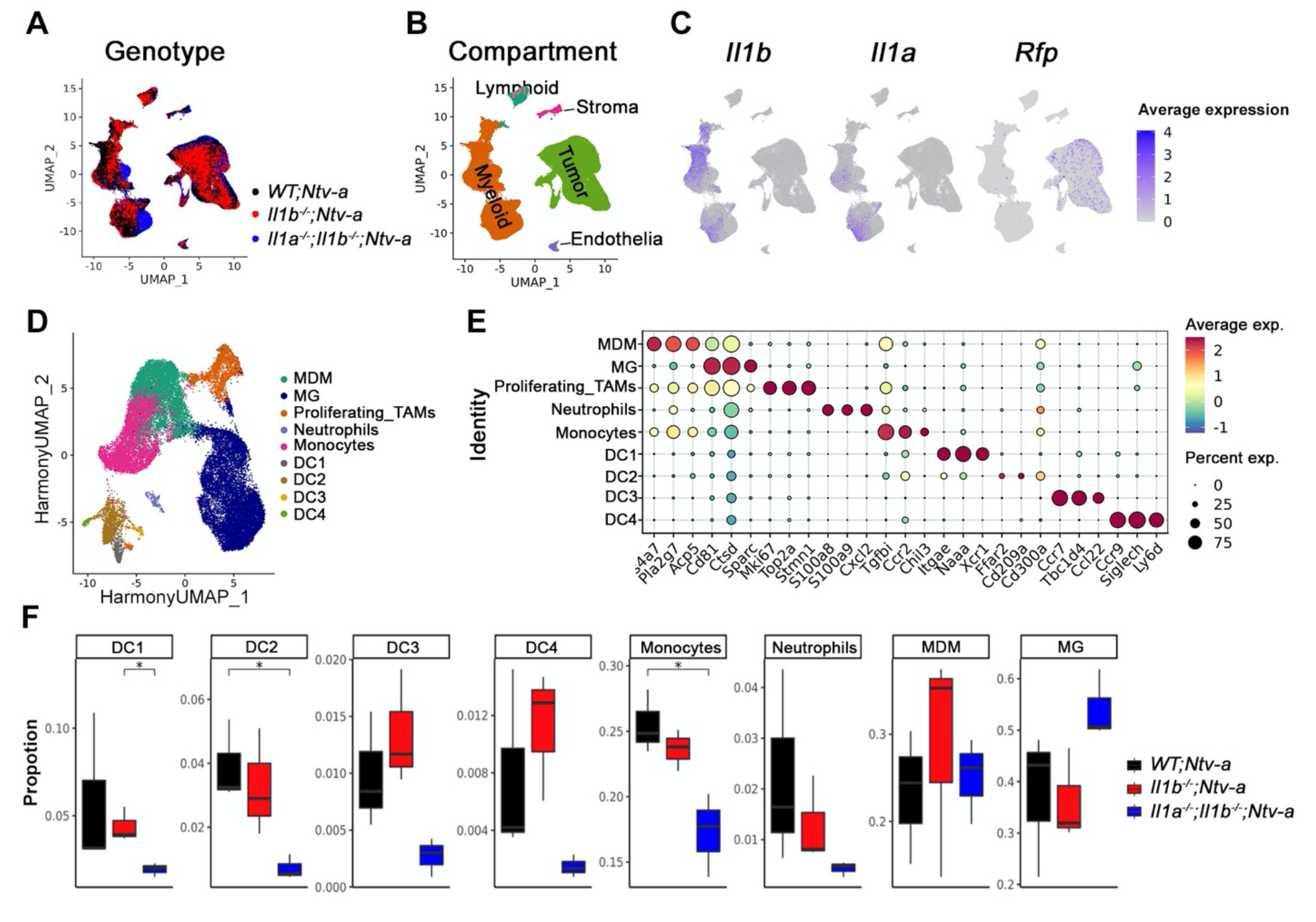
scRNA-seq reveals reduction in inflammatory monocytes in *Il1b^-/-^;Ntv-a* mice. (**A**) UMAP dimensionality reduction of the scRNA-seq data isolated from PDGFB-driven tumors *WT;Ntv-a* (*black*), *Il1b^-/-^*;*Ntv-a* (*red*) and *Il1a^-^;Il1b^-/-^;Ntv-a* (blue) mice. (**B**) UMAP of single cells in A colored by annotated cell class. (**C**) *Il1a*, *Il1b*, *RFP* expression overlayed on the same UMAP coordinates. (**D**) UMAP dimensionality reduction of myeloid cells in the tumors colored by annotated myeloid subtypes. (**E**) Dot plot of selected marker genes defining myeloid sub types. (**F**) Composition of myeloid cell subpopulations in tumors generated in *WT;Ntv-a, Il1b^-/-^;Ntv-a* and *Il1a^-^;Il1b^-/-^;Ntv-a* mice. Two-tailed Student’s *t*-test, *P<0.05.

Next, we stratified the myeloid cells into phenotypical and functional subgroups (**Fig. 4D**), with *post-hoc* annotation based on their expression profiles of signature genes (**Fig. 4E**). We identified two tumor-associated macrophage populations based on their ontogeny, namely monocyte-derived macrophages (MDM) and MG (**Fig. 4E**). We found that the most significant changes occurred in the number of monocytes and DC2 dendritic cells, while other cell types varied only marginally between these three genotypes (**Fig. 4F**). Specifically, we found a reduction in the relative abundance of monocytes in *Il1b;Ntv-a* compared to *WT;Ntv-a* mice, which was more pronounced in the *Il1a^-/-^;Il1b^-/-^;Ntv-a* double KO mice (**Fig. 4F**).

To complement and corroborate the scRNA-seq data, we used multi-parameter spectral flow cytometry (**Fig. 5**) to distinguish between bone marrow-derived inflammatory monocytes (CD11b^+^CD45^Hi^Ly6c^Hi^Ly6g^Neg^CD49d^+^), monocyte-derived macrophages (MDM, CD11b^+^CD45^H1^Ly6c^Lo/Neg^Ly6g^Neg^CD49d^+^), microglia (CD11b^+^CD45^Lo^Ly6c^Neg^Ly6g^Neg^CD49d^Neg^), and neutrophils (CD11b^+^CD45^+^Ly6c^+^Ly6g^+^CD49d^+^) (**Fig. 5A**, gating strategy shown in **Fig. S3**). There was a significant reduction in total myeloid cells in tumors from *Il1b^-/-^;Ntv-a* mice relative to *WT;Ntv-a* mice (**Fig. 5B**, P<0.01), which was confirmed by IBA1 immunohistochemistry (**Fig. S4**, P<0.01). Among these CD11b^+^myeloid cells, microglial abundance was decreased in *Il1b^-/-^;Ntv-a* mice, but failed to reach statistical significance (**Fig. 5C**, P=0.1667). However, the abundance of infiltrating BMDMs was reduced in *Il1b^-/-^;Ntv-a* mice (**Fig. 5C**, P<0.05). Using FlowSOM (**Fig. 5D**) to identify subpopulations within the BMDM compartment (gating strategy illustrated in **Figure 5E**), we found a significant decrease in Ly6c^Hi^ (newly infiltrated inflammatory monocytes) in *Il1b^-/-^;Ntv-a* mice, without any change in Ly6c^Lo/Neg^(differentiated macrophages) and Ly6g^+^neutrophils (**Fig. 5F**), closely reflecting the scRNA-seq data (**Fig. 4F**). Similarly, we detected a reduction in CD24^+^CD103^-^ DC2 cells (**Fig. 5G**) in tumors generated in *Il1b^-/-^;Ntv-a* mice compared to *WT;Ntv-a* mice, with no change in DC1 cells (**Fig. S5**).

**Figure 5.**
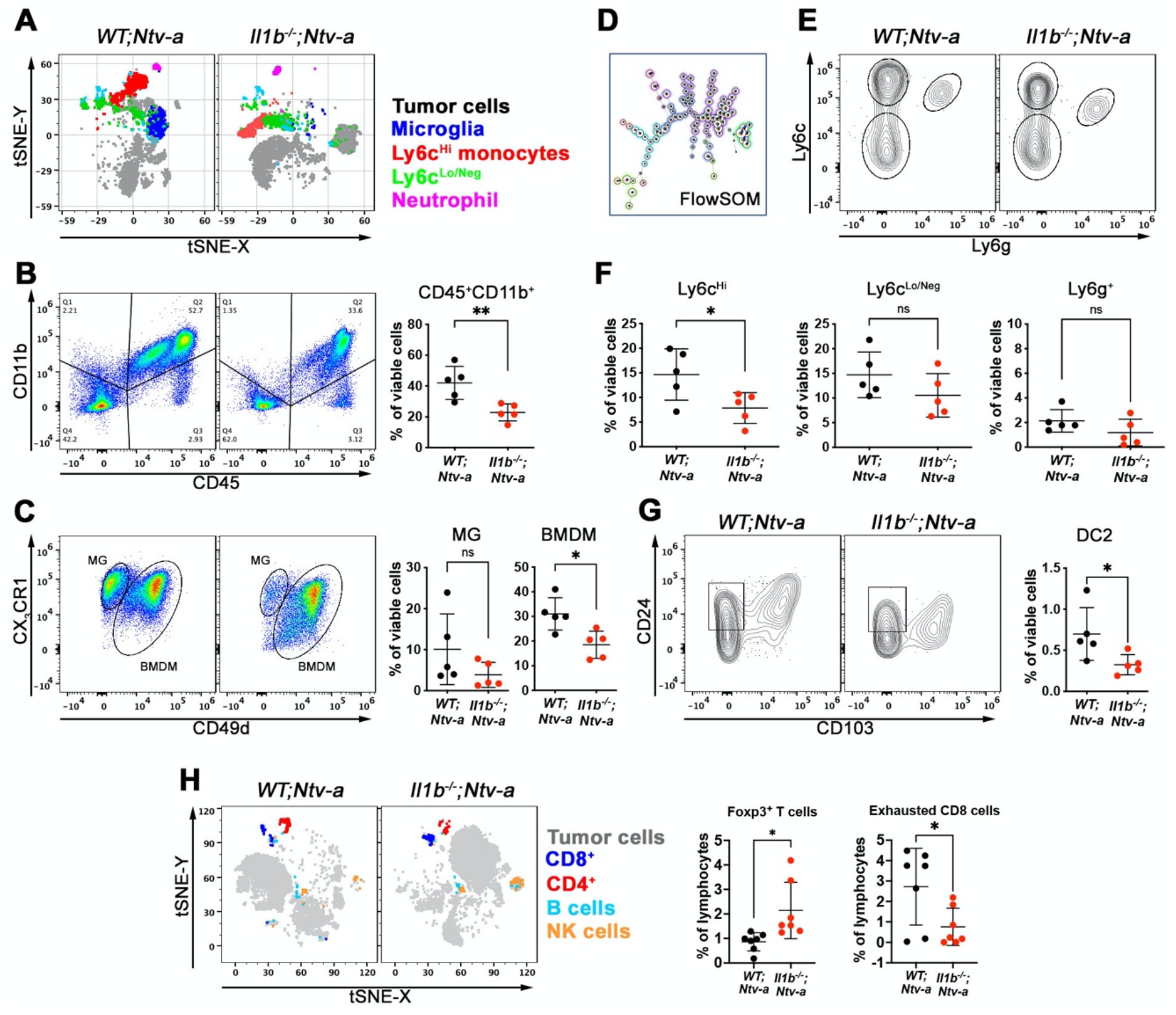
*Il1b* ablation reduces the influx of inflammatory monocytes and DC2 cells into tumors of mice with *PDGFB-driven* GBM. (**A**) tSNE plots of spectral flow cytometry illustrating the tumor cell/lymphoid composition in *WT;Ntv-a* an *Il1b^-/-^;Ntv-a* mice bearing *PDGFB*-driven GBM. (**B**) and (**C**) illustrate the gating strategy for myeloid cells, discriminating between resident brain microglia and BMDM, with corresponding quantification dot graphs. Two-tailed Student’s *t*-test, *P<0.05, **P<0.01 (**D**) Illustration of FlowSOM differentiating subtypes of bone marrow derived myeloid cells. (**E**) Representative contour plots for gating monocytes (Ly6c^hi^), differentiated macrophages (Ly6c^low/neg^), and neutrophils (Ly6g^+^). (**F**) Quantification of myeloid cell subtypes depicted in **E**. Two-tailed Student’s *t*-test, *P<0.05, ns=not significant. (**G**) Representative plots and quantification of DC2 populations in *WT;Ntv-a* and *Il1b^-/-^;Ntv-a* mice bearing *PDGFB*-driven GBM. Two-tailed Student’s *t*-test, *P<0.05. (**H**) tSNE plots and quantification of lymphoid cells by spectral flow cytometry. Two-tailed Student’s *t*-test, *P<0.05.

To determine whether this reduction in Ly6c^Hi^ monocytes in the tumors reflected fewer Ly6c^Hi^ monocytes in the circulation, we enumerated cell numbers in whole blood of mice using flow cytometry. We found no difference in the myeloid cells between tumor-bearing *WT;Ntv-a* and *Il1b^-/-^;Ntv-a* mice (**Fig. S6**). Together, these data demonstrate that IL-1 β deficiency selectively reduces monocyte infiltration to the tumor from the periphery.

Next, we performed in-depth analysis on the lymphocyte compartment by scRNA-seq (**Fig. S7**) and spectral flow cytometry (**Fig. S8**). Lymphoid cells account for a small portion of infiltrating cells in GBM, consistent with numerous previous studies (Chen and Hambardzumyan, 2018; Han et al., 2016). We found a slight increase of Foxp3^+^ T_reg_ by both scRNA-seq and flow cytometry (**Fig. S7** and **Fig. 5H**, P<0.05), and a significant decrease in the amount of exhausted (Tim3^+^PD-1^+^) CD8 T cells (**Fig. 5H**, P<0.0), but not in other cell types (**Fig. S9**). Although with a small footprint in the tumor, this reduction in exhausted T cells may be an important contributor to the prolonged survival of the *Il1b^-/-^;Ntv-a* mice.

### IL-1β increases monocyte recruitment via tumor cell expression of MCPs

As shown above, reduced inflammatory monocyte infiltration is one of the most prominent features observed in tumors generated in *Il1b^-/-^;Ntv-a* mice. This effect closely aligns with our previous findings demonstrating increased *Il1b* expression in tumor-bearing *Cx3cr1^-/-^* mice, which have increased inflammatory monocytes and shortened survival times (Feng et al., 2015). We have previously shown that primary *PDGFB*-driven tumors increase CCL2 expression in response to recombinant IL-1β (rIL-1β) (Feng et al., 2015). However, genetic *Ccl2* deletion did not reduce TAM content in *PDGFB*-driven tumors (Chen et al., 2017).

In contrast to *Ccl2, Il1b* loss was sufficient to reduce both inflammatory monocytes and total TAM numbers in *PDGFB*-driven tumors, suggesting that other MCP family members, like CCL7, CCL8, and CCL12 (or CCL13 in humans), could be essential for mediating inflammatory monocyte GBM infiltration. To test this hypothesis, we analyzed IL-1β induction of all four MCP members. First, we found that all MCP members were increased in primary *PDGFB*-driven mGBM relative to normal brain, and all but one (CCL8) was increased in primary *Nf1*-silenced mGBM (**Fig. S10A**, 2 to 5 times log_2_ fold change, P<0.05). Next, we examined this induction under two different culture conditions for (1) enriched glioma stem cells (GSCs) grown in a serum-free neural stem cell medium supplemented with EGF and β–FGF (termed neurosphere condition-NSC), and (2) bulk tumor cells grown in 10% FBS in DMEM medium (termed differentiation condition, or FBS) (Feng et al., 2015) (**Fig. S10B**). We found that freshly-sorted *PDGFB*-tumor cells expressed higher levels of CCL2, CCL7, CCL8, and CCL12 relative to the same cells maintained *in vitro* (up to 6 passages) under both GSC and FBS conditions (**Fig. S10C**, P<0.05). Although *Ccl12* mRNA levels did not reach statistical significance, it was higher in fresh isolates than in cultured cells (**Fig. S10C**). These observations suggest that stromal support might be essential for *PDGFB*-driven tumor cells to produce MCPs.

To examine this potential explanation, we grew freshly-isolated *PDGFB*-driven tumor cells under NSC and FBS conditions, and measured MCP levels in the supernatant by ELISA at passage 1 (P1). While MCPs were expressed under both NSC and FBS conditions at P1 (**Fig. S10D**), at later passage (P6), *PDGFB*-driven primary cells had reduced MCP expression, while *Nf1*-silenced cells showed high expression regardless of the number of passages (**Fig. S10E**), indicating that *Nf1*-silenced tumor cells are less reliant on stromal support for MCP induction, which is consistent with the high levels of IL-1β in the tumor cells of this GBM molecular subtype (MES).

Based on these observations, we next hypothesized that supplementing stroma-derived factors, such as IL-1β, could restore *PDGFB*-driven (PN), but not *Nf1*-silenced (MES), tumor cells production of MCPs. As such, we added IL-1β (100 pM) to *PDGFB*-driven primary and *Nf1*-silenced tumor cell cultures under both NSC and FBS conditions. While IL-1β stimulation had no effect on RNA or protein expression in *Nf1*-silenced tumor cells (**Fig. S11A, B**), MCP RNA and protein expression levels increased in *PDGFB*-driven tumor cells under both NSC and FBS conditions (**Fig. S11C, D**). Together, these data support that stroma-derived IL-1 β is essential for mediating monocyte infiltration through the induction of MCP production in *PDGFB-* driven GBM cells.

### IL-1β activates NF-κB signaling in tumor cells in a GBM subtype-specific manner

Based on prior studies implicating the NF-*κ*B pathway in IL-1β-mediated signal transduction, we analyzed this pathway in *PDGFB*-driven and *Nf1*-silenced GBM tumor cell cultures *in vitro*. For *PDGFB*-driven tumors, we used primary cells freshly isolated from excised tumors (P1 to P3). For *Nf1-* silenced tumors, we used murine GBM lines (1816 and 4622), which lack *Nf1* and *Tp53* expression (Gursel et al., 2011; Pan et al., 2017; Reilly et al., 2000). We chose these lines because *Nf1*-silenced GBM cell are difficult to maintain *in vitro*. *Nf1*-silenced tumor cells have increased phosphorylation (activation) of NF-*κ*B pathway intermediates relative to *PDGFB*-driven cells (**Fig. 6A**), consistent with a recent study showing that MES GBM GSCs exhibit constitutively active NF-*κ*B signaling (Bhat et al., 2013). We next evaluated the effect of IL-1β (100 pM) on NF-*κ*B pathway activation in NSC or FBS *PDGFB*-driven and *Nf1*-silenced tumor cell cultures. While IL-1β induced NF-*κ*B activation under both FBS (**Fig. 6B**, P<0.05) and NSC (**Fig. S12**, P<0.05) conditions in *PDGFB*-driven tumors, IL-1β stimulation did not induce NF-*κ*B pathway activation in *Nf1*-silenced tumor cells (**Fig. 6C**). These results indicate that *PDGFB*-driven and *Nf1*-silenced tumors intrinsically differ in NF-*κ*B pathway signaling and responses to IL-1β stimulation. While *PDGFB*-driven tumors secrete MCPs in response to IL-1β, *Nf1-* silenced tumors reach a plateau of MCP production, even without IL-1β stimulation, due to constitutively active NF-*κ*B pathway signaling.

**Figure 6.**
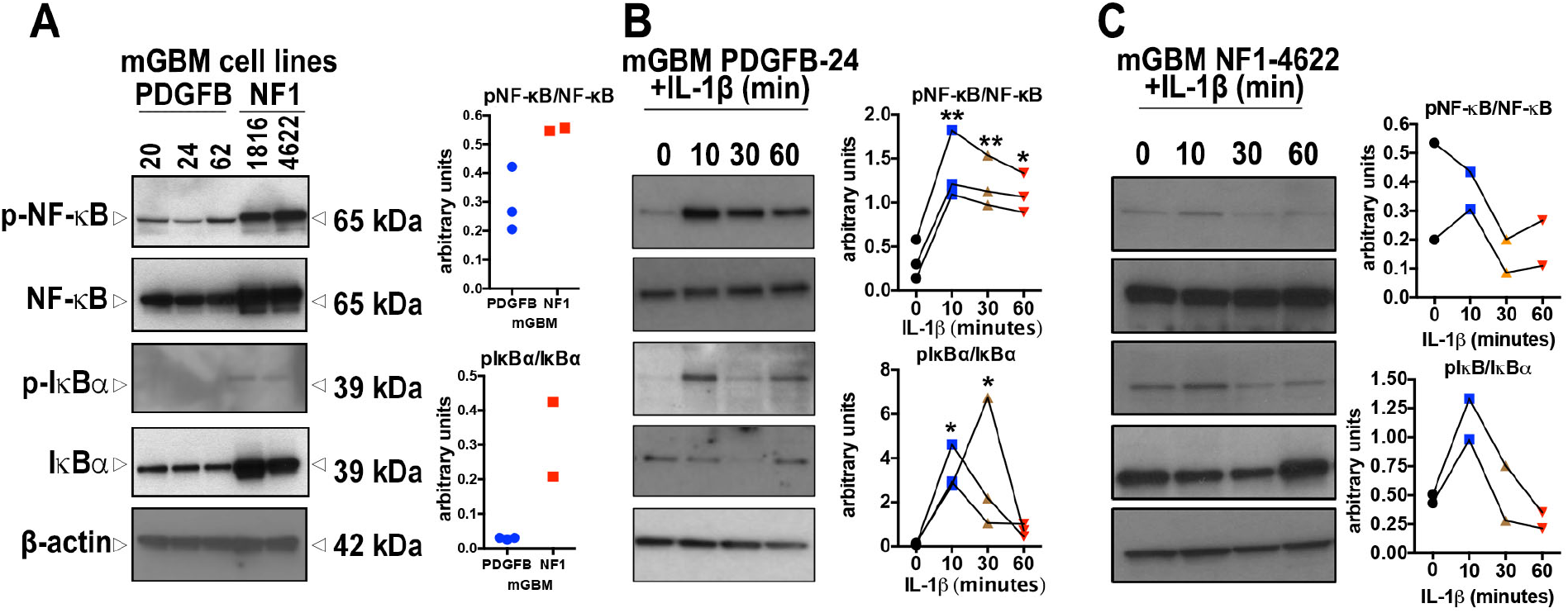
IL-1β induces NF-κB pathway activation in PDGFB-driven, but not *Nf1-* silenced, GBM cell cultures. (**A**) Immunoblot showing NF-κB pathway components in both *PDGFB-* driven and *Nf1*-silenced mGBM cultures *in vitro*.(**B**) NF-κB pathway components in response to IL-1β treatment of *PDGFB-* driven GBM cells in FBS-containing medium, quantification with triplicate experiments. *P<0.05, **P<0.01 by ANOVA test. (**C**) NF-κB pathway components in response to IL-1β treatment of *Nf1*-silenced GBM cells in FBS-containing medium and quantification with triplicate experiments.

To confirm that this effect was not an *in vitro* cell culture-specific phenomenon, we generated *PDGFB*-driven tumors in *WT;Ntv-a, Il1b^-/-^;Ntv-a*, and *Il1a^-/-^;Il1b^-/-^;Ntv-a* mice for ELISA determinations of CCL2, CCL7, CCL8 and CCL12 expression. Loss of *Il1* leads to decreased MCP production, which was more pronounced in tumors generated in *Il1a^-/-^;Il1b^-/-^;Ntv-a* mice compared to *Il1b^-/-^;Ntv-a* mice (**Fig. S13**). The reduction in MCP in *Il1b^-/-^* tumors is not apparent relative to *WT* mice, which might reflect the fact that IL-1β-producing BMDMs are restricted to perinecrotic/perivascular areas of tumors and the ELISA was performed on whole tumor lysates. Furthermore, because *Il1a* expression is more wide-spread in the tumors, the effect on MCP reduction might be larger from the combined loss of both *Il1a* and *Il1b* in double KO mice. Finally, we observed no differences in VEGFA levels, which was used as an internal control (**Fig. S13**, P=0.63). Taken together, these data establish that loss of IL-1 in the tumor and stroma leads to a reduction in MCPs *in vivo*, and this effect correlates with decreased infiltration of inflammatory monocytes into *PDGFB*-driven tumors (**Fig. 4F**).

### Germline *Il1a and Il1b* loss reverses the survival benefit conferred by *Il1b*-deletion

In light of the strong effect of combined *Il1a* and *Il1b* loss on MCP levels in tumors, we sought to determine whether genetic deletion of both *Il1a* and *Il1b* would further extend survival of tumor-bearing mice. Surprisingly, in contrast to the extended survival duration of *Il1b^-/-^;Ntv-a* mice (**Fig. 3A**), the survival time of tumor-bearing *Il1a^-/-^;Il1b^-/-^;Ntv-a* mice was similar to that of the *WT;Ntv-a* mice (**Fig. 7A**). Consistently, we did not observe any differences in the number of phosphor-H3-positive proliferating cells in the tumors between these two genotypes (**Fig. S14**).

**Figure 7.**
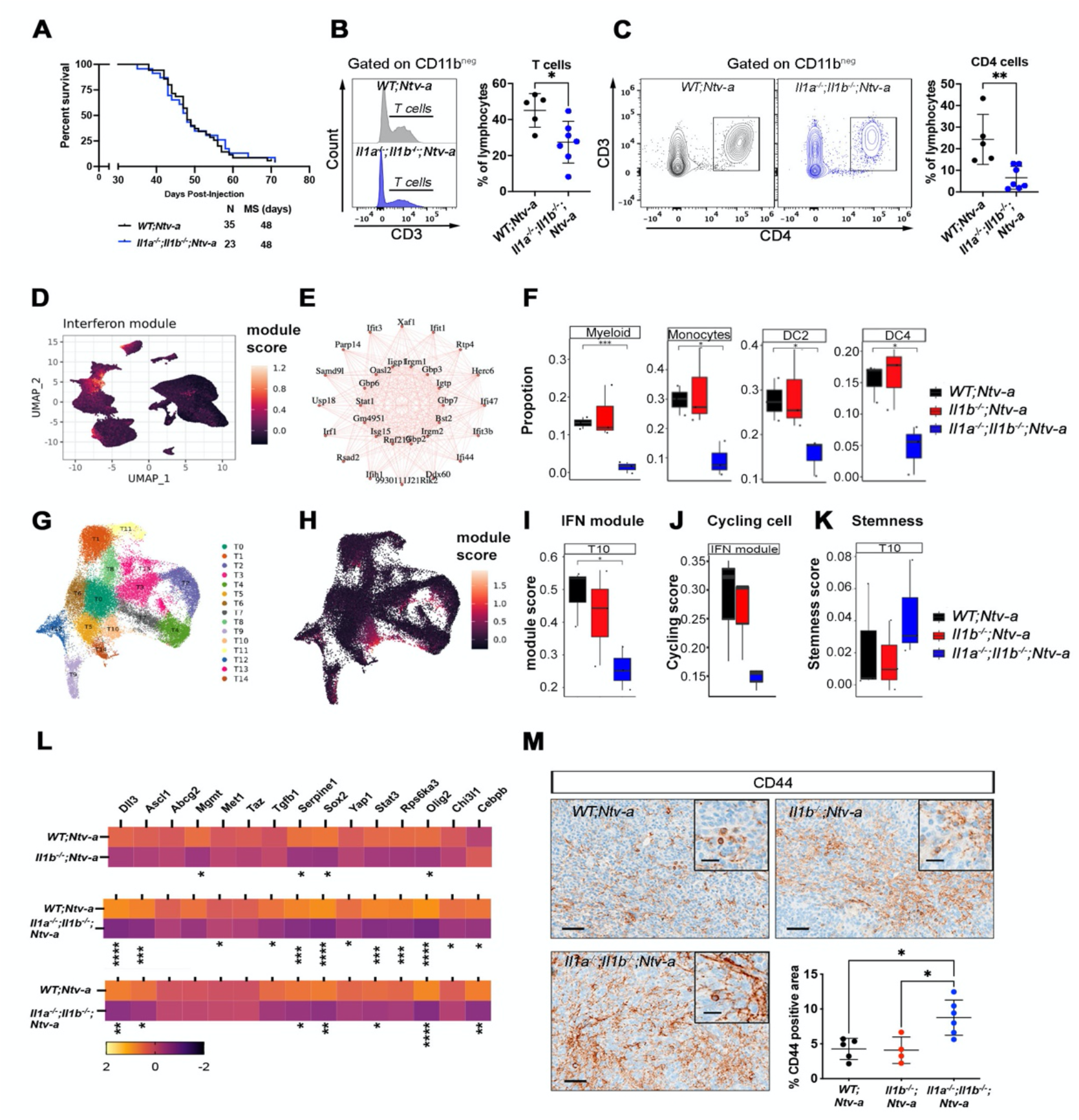
Genetic ablation of *Il1a/b* has no impact on the survival of *PDGFB-driven* GBM-bearing mice. (**A**) Kaplan-Meier survival curves of *PDGFB*-driven tumors generated in *WT;Ntv-a* and *Il1a^-/-^;Il1b^-/-^;Ntv-a* mice. (**B**) Histogram and quantification of CD3 T cells examined by spectral flow cytometry. Two-tailed Student’s *t*-test, *P<0.05. (**C**) Dot plots of CD4 T cells and its quantification examined by spectral flow cytometry. Two-tailed Student’s *t*-test, **P<0.01. (**D**) UMAP dimensionality reduction of all cells examined by scRNA-seq and colored by expression of the IFN module derived by WGCNA. (**E**) Interconnected graph showing the top 30 most co-expressed genes in the IFN module. (**F**) Proportion of myeloid cell types highly expressing the interferon module for each genotype. Two-tailed Student’s *t-* test, *P<0.05. (**G**) UMAP dimensionality reduction of tumor cells, colored by assignment to 15 tumor clusters. (**H**) IFN module score overlayed on the same UMAP coordinates. (**I**) IFN module score of cluster T10. (**J**) Proportion of proliferating malignant cells found in cluster T10, grouped by genotype. (**K**) Quiescent score of malignant cells found in cluster T10. (**L**) qPCR analysis of expression of genes associated with stemness signatures in tumors generated in *WT;Ntv-a, Il1b^-/-^;Ntv-a andIl1a^-/-^;Il1b^-/-^;Ntv-a* mice. Two-tailed Student’s *t*-test, *P<0.05, **P<0.01; ***P<0.001; ****P<0.0001. (**M**) Immunohistochemistry analysis of CD44 with quantification. *P<0.05 by one-way ANOVA with Tukey’s *post-hoc* test.

To define the etiology for this paradoxical result, we analyzed immune cell content in these tumors by flow cytometry, and found a reduction in the abundance of total myeloid cells, lymphoid cells, MG and BMDM in *Il1a^-/-^;Il1b^-/-^;Ntv-a* mice (**Fig. S15A**, P<0.05). This decreased MG abundance, which was not apparent in *Il1b^-/-^;Ntv-a* tumors (**Fig. 5C**, P=0.1667), might reflect decreased proliferation due to the loss of IL-1 signaling (Bruttger et al., 2015). In contrast to *Il1b^-/-^;Ntv-a* mice, tumors generated in *Il1a^-/-^;Il1b^-/-^;Ntv-a* mice showed reduced CD3^+^ T cell infiltration (**Fig. 7B**, P<0.05), which resulted mainly from fewer CD4 T helper cells (**Fig. 7C**, P<0.01). This reduction in both MG and CD4 T cells may, at least partially, explain the survival differences seen between in *Il1b^-/-^;Ntv-a* and *Il1a^-/-^;Il1b^-/-^;Ntv-a* mice; however, other factors may also contribute to the shortened survival of *Il1* KO mice.

To define the mechanism(s) underlying the effects seen in *Il1a^-/-^;Il1b^-/-^;Ntv-a* mice, we performed weighted gene co-expression network analysis (WGCNA) adopted for scRNA-seq data. This analysis reveals gene regulatory network, or “modules”, based on gene co-expression patterns (Langfelder and Horvath, 2008). This analysis enabled us to identify gene expression modules shared across cell types that would not be apparent using Seurat’s Louvain clustering method. Among all the modules, the IFN module (**Fig. 7D**) demonstrated the most significant difference between the three genotypes. The top 30 most co-expressed genes in this module are shown in an interconnected graph (**Fig. 7E**). We found a substantial decrease of the interferon (IFN) signaling pathway in the double KO mice across many cell types, including total myeloid cells (P<0.001), monocytes (P<0.05), DC2 (P<0.05), and DC4 cells (P<0.05, **Fig. 7F**). Similarly, unsupervised clustering on malignant cells identified 15 sub-clusters (**Fig. 7G** and **Fig. S16**), where cells bundled in cluster 10 (T10) exhibited impaired IFN signaling responses specifically in *Il1a^-/-^;Il1b^-/-^;Ntv-a* mice (**Fig. 7H&I**). Quite unexpectedly, when we examined the proliferation properties of these cells by cell cycle analysis, we found that almost all the T10 cells in *Il1a^-/-^;Il1b^-/-^;Ntv-a* mice were not proliferating, suggesting they may be in a quiescent state (**Fig. 7J**). This observation inspired us to speculate that these cells could be more stem-like cells entering quiescence in response to an absence of IFN stimulation from the myeloid cells. Consistent with this idea, scRNA-seq analysis revealed an upregulation of markers associated with quiescent stem-like GBM cells in cluster T10 (**Fig. 7K**) (Mukherjee, 2020).

Using a previously established quantitative PCR (qPCR) panel to examine the molecular profiles of stem-like cells in mGBM (Herting et al., 2017), we analyzed the tumors generated in *WT;Ntv-a*, *Il1b^-/-^;Ntv-a* and *Il1a^-/-^;Il1b^-/-^;Ntv-a* mice. Although some of these genes remained unchanged between the three genotypes, we observed an IL-1 dosedependent reduction (stronger effects when both *Il1a* and *Il1b* are lost relative to *Il1b* loss alone) in *Sox2*, *Olig2*, and *Serpine1* expression (Gimple et al., 2019) (**Fig. 7L**). In contrast to *Il1b*-null tumors, tumors from combined *Il1a/b* loss mice showed a reduction *in Ascl1* expression levels, important for the establishment of neuronal fate and loss of selfrenewal (Park et al., 2017). Loss of *ASCL1* also inversely correlates with the expression of CD44, another prominent stem cell marker (Pietras et al., 2014), which we confirmed by immunohistochemistry (**Fig. 7M**, P<0.05). Taken together, these finding indicate that cancer progression is promoted by combined loss of both IL-1a and IL-1β.

### Local antagonism of IL-1β or IL-1R1 prolongs the survival of tumor-bearing mice

Since all of the genetic experiments were performed in germline *Il1a/b*- or *Il1b*-deficient mice, we sought to determine whether locally neutralizing IL-1β or introducing an IL-1R1 antagonist (IL-1Ra) would impede tumor growth and prolong the survival of tumor-bearing mice, without interfering with normal development and the function of immune cells. For these experiments, we created *PDGFB-* driven GBM in *WT;Ntv-a;* mice (or *WT;Ntv-a;Cdkn2a^-/-^* mice, and observed the similar results between these two strains) and allowed the tumors to grow for 15 days. At this time, we installed guide cannulas into the brains of each mouse through the same burr hole that was used for tumor initiation (**Fig. 8A**). These cannulas enabled us to repeatedly deliver anti-IL-1β antibodies or IL-1Ra on a daily basis without damaging brain tissue. This direct delivery system also avoided the caveat of low bioavailability when the compounds are injected via the intraperitoneal route. We divided the mice into three groups, receiving either vehicle control, IL-1Ra, or anti-IL-1β antibody daily. Kaplan-Meier survival curves demonstrate that pharmacologically antagonizing IL-1β or IL-1R1 prolonged the survival of GBM-bearing mice relative to vehicle control (**Fig. 8B**, P<0.05). Interestingly, when we examined the tumor tissues of these mice with IHC (**Fig. 8C**), we found a reduction in IBA1-positive cell density, suggesting a reduction in TAM infiltration (**Fig. 8D**, P<0.05). In fact, reduced TAM content in the tumor serves as an indicator of improved prognosis (**Fig. 8E**). Together, these results establish a rationale for clinical translation of local administration of IL1-targeted therapy for GBM.

**Figure 8.**
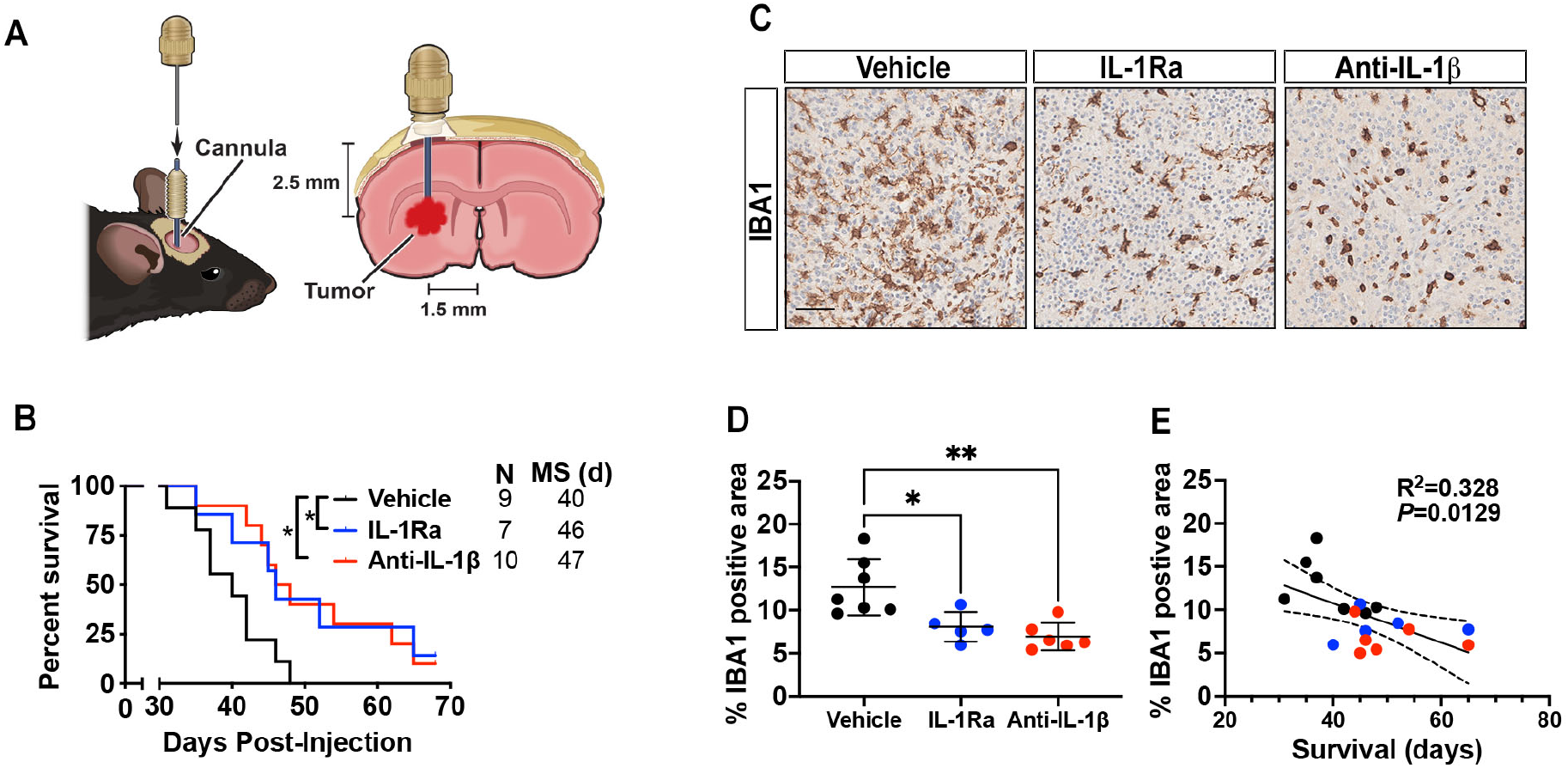
Intra-tumoral anti-IL-1β antibody or IL-1Ra administration prolongs survival of *PDGFB-driven* GBM-bearing mice. (**A**) Illustration of the experimental design. (**B**) Kaplan-Meier survival curves of *PDGFB-driven* tumor-bearing *Ntv-a;Cdkn2a^-/-^* mice treated following treatment with vehicle, anti-IL-1β Ab, or IL1R antagonist. *P<0.05 by MC test. (**C**) Representative IBA1 IHC images of tumors from vehicle-, anti-Il-1B antibody or IL1R antagonist-treated tumor-bearing mice. (**D**) Quantification of IBA1-positive tumoral areas. One-way ANOVA, Tukey’s multiple comparison test, *P<0.05.**P<0.01. (**E**) IBA1 expression correlates with survival of tumor-bearing mice. Color of data points match the genotypes depicted in D.

## Discussion

In this study, we analyzed the mechanistic role of IL-1 signaling in driving TAM-tumor cell cross-talk that promotes tumor progression. Using both gold-standard genetic and pharmacological tools, we demonstrated that local suppression of IL-1 β/IL-1R1 signaling can significantly prolong the survival of GBM-bearing mice. Previous studies investigating the role of IL-1 signaling, specifically that of IL-1β, in GBM cell cultures *in vitro* have yielded inconsistent results: some studies show that IL-1β can inhibit tumor growth (Oelmann et al., 1997), while others demonstrate that it promotes tumor development, invasion, and cancer stem cell frequency in patient-derived primary GBM cell lines (Feng et al., 2015; Kai et al., 2021; Sarkar and Yong, 2009). Further, studies exist showing that there is no impact of IL-1β on tumor cell proliferation (Sharma et al., 2011a). The complexity of these observations is further compounded due to many of these studies using the same glioma cell line but documenting different results. Based on the duration of rIL-1β treatment, it has been shown that short-term treatment led to growthpromoting effects (Lachman et al., 1987), while longer rIL-1β treatment (30 days) resulted in growth suppression (Tanaka et al., 1994). Overall, these published results indicate that the effects of IL-1β on GBM cells are cell line-specific, culture condition-determined, and time-dependent; thus, signifying that any *in vitro* investigations may not reliably recapitulate the complexity IL-1b function in pathophysiological conditions *in vivo*. Here, we set out to determine whether tumor genotype and/or timing of IL-1β inhibition (germline loss versus local tumor targeting) impact GBM growth *in vivo* by using GEMMs. We also investigated whether there are differences in the effects of individual IL-1 ligands (IL-1α vs IL-1β) on GBM growth and myeloid cell recruitment into the tumors. Our results demonstrate that both RNA and protein levels of *IL1B* are increased in *IDH-WT* GBM patient samples, and this elevated expression of *IL1B* inversely correlates with patient survival. Genetic deletion of *Il1b*, or locally suppressing IL-1 signaling within the tumors, can significantly prolong the survival of GBM-bearing mice. Our results have provided rationale for using IL-1 β-targeted therapy as a practical and effective treatment for GBM.

We observed a significantly higher level of IL-1 β in human MES GBM than PN GBM, at both the RNA and protein levels (**Fig. 2**). This difference remains true in their corresponding murine models, with *Nf1*-silenced mGBM demonstrating elevated IL-1 β as compared to *PDGFB*-driven mGBM. This observation should not come as surprise though, as it has been established MES GBM are more “inflamed” and immuno-active than PN GBM (Kaffes et al., 2019). However, the novel observation we made in this study is that MES GBM cells have constitutive activation of NF-κB pathway, which can explain its heightened immunological reactions. This observation is in agreement with a previous study demonstrating active NF-κB signaling in human MES, but not PN, GSC cultures (Bhat et al., 2013). These results clearly illustrate that the effect of IL-1β is genotype-specific and can partially explain the discrepancies in the literature where tumor cell genotype was not considered as a covariate. It appears that, at least *in vitro*, MES cells may have reached a plateau in NF-κb activity, thereby any exogenous IL-1β supplemented to the cell culture did not further increase the phosphorylation of NF-κB (**Fig. 6**). It has been documented *Il1b* expression can be regulated by NF-κB (PMID: 8413223), which can explain increased levels of IL-1β in NF1-silenced tumors and primary cultures.

Decreased inflammatory monocyte recruitment in *Il1b*-deficient mice was similar to what we observed previously in *Il1r1*-deficient mice (Herting et al., 2019), which we hypothesized was due to diminished MCP production and thereby, reduced monocyte recruitment via chemotaxis. The effect of *Il1* ablation on the MCP network *in vivo* was more pronounced when both *Il1* ligands were ablated. Interestingly, in tumors lacking the *Il1b* gene, there was a significant induction of chemokine CCL7, which was in contrast to the *in vitro* data performed with tumor cells cultures (**Fig. S13**). This increase in CCL7 *in vivo* could possibly be due to its increased production by other cell types in the TME, similar to what was shown in a lung cancer model, where loss of CCL7 accelerated tumor growth due to impaired function of the antitumor immune response (Zhang et al., 2020).

Ablation of both *Il1a* and *Il1b in vivo* resulted in more potent decreases in both monocyte recruitment and total TAMs in the tumors, however, to our surprise, the survival duration did not differ from that of the *WT;Ntv-a* mice. Unexpectedly, our scRNA-seq data revealed that there is a significant disruption of the IFN signaling pathway in both the myeloid compartment and malignant cells in *Il1a^-/-^;Il1b^-/-^;Ntv-a* double KO mice. Although playing divergent roles in host defense, it is known that IL-1 and IFN pathways can cross talk with each other in various diseases (Mayer-Barber and Yan, 2017). Type I IFN can actively regulate IL-1 expression and may also antagonize its biological functions; however, much less is known about how IL-1 could in turn regulate IFN production and/or its effector functions (Mayer-Barber and Yan, 2017). Here, we demonstrated that IFN function is impaired in *Il1* KO mice in mGBM. Interestingly, IFNs of both type I and type II have been shown to be crucial in cancer-immunoediting process where they are considered central coordinators of tumor-immune interactions, and play a pivotal role in promoting antitumor responses (Dunn et al., 2006). When IFN functions are impaired, tumors in *Il1a/b*-KO mice demonstrated increased stemness features, thereby raise their aggressiveness (**Fig. 7**). Based on this observation, we conclude that complete ablation of *Il1* results in the impairment of IFN signaling in GBM, which escapes the immune control of malignancy and leads to earlier demise of the mice.

Pharmacological antagonizing *Il1* signaling locally in tumor-bearing mice extended survival time and decreased TAM infiltration, opposite to what was observed with *Il1a^-/-^;Il1b^-/-^;Ntv-a* mice. This observation suggests the differences seen in *Il1*-deleted mice can be attributed to germline loss of *Il1* that leads to decreased lymphoid infiltration and the loss IFN control of malignancy, although in these mice TAMs infiltration is also diminished. Therefore, reduction in tumor-promoting TAM alone cannot offset the combined detrimental effects of lymphoid and IFN loss in *Il1a^-/-^;Il1b^-/-^;Ntv-a* mice. On the contrary, local delivery of IL-1-targeted therapy in immunocompetent mice reduced TAM infiltration but did not alter IFN signaling, thereby providing effective inhibition of tumor growth. Our results also indicate that IL-1α and IL-1β may perform distinct functions in GBM. Several mechanisms account for these differences. First, IL-1α can be produced by nearly all cells and is typically released upon cell death; while IL-1β is mainly produced in activated myeloid cells as a pro-protein, which is proteolytically processed to its active form by caspase 1 following inflammasome formation and activation, leading to mature IL-1 β release (Barksby et al., 2007; Chen et al., 2017; Dinarello et al., 2012; March et al., 1985). Secondly, unlike IL-1β which functions exclusively through binding IL-1R1 on the plasma membrane, IL-1α can translocate into the nucleus where it binds transcription factors and activates expression of proinflammatory cytokines and chemokines independently of IL-1R1 signaling (Di Paolo and Shayakhmetov, 2016). Interestingly, IL-1α can serve as a double-edged sword during tumor development (Malik and Kanneganti, 2018). In certain cancers such as fibrosacoma, overexpression of IL-1α leads to regression of the disease via enhanced immune-surveIl1ance mediated by CD8^+^ T cells, NK cells and M1 macrophages (Dvorkin et al., 2006; Voronov et al., 1999).

Ly6c^Hi^Ly6g^Neg^ cells are also referred to as monocytic myeloid-derived suppressor cells (mMDSCs), whereas the Ly6c^Hi^Ly6g^+^ cells are termed granulocytic myeloid-derived suppressor cells (gMDSCs). A recent study showed a dichotomous distribution of these two suppressor cell types in different genders of GBM-bearing mice when the tumors were generated by transplanting GL261 tumors isolated from a male donor (Bayik et al., 2020). Here, however, we did not observe any differences in mMDSC and gMDSC infiltration when we stratified our data by gender (data not shown). Additionally, to evaluate whether the gender effect observed in the previous publication were a donor-sex specific result, we performed experiments where male and female donor-derived primary tumor cells were transplanted into male and female recipients, as illustrated in **SFig17**. We did not observe gender-specific differences in their myeloid composition or the survival time of tumor-bearing mice (**Fig. S17B and C**). It is possible that the differences between our current data and the prior report are model-specific and therefore warrants further investigation.

In summary, our findings demonstrate a pro-tumorigenic function of IL-1β in mGBM, and blocking IL-1β signaling decreases inflammatory monocyte recruitment, microglia abundance, and the frequency of exhausted CD8+ T cells, which ultimately prolongs the survival of tumor-bearing mice. Along with our previous findings showing targeting IL-1β can effectively reduce cerebral edema in GBM patients (Herting et al., 2019), our results here further support the application of antagonism of IL-1β as a promising therapy for GBM. Together, these studies provide strong rationales for clinical translation of antagonizing IL-1β in treating GBM.

## Acknowledgements

We would like to acknowledge Emory Children’s Flow Cytometry Core and the Integrated Cellular Imaging Cores for their services. We would also like to acknowledge the Mount Sinai Flow Cytometry Core, as well as the Genomics Core for scRNA-seq services. We extend our thanks to Mr. David R. Schumick for generating illustrations and Dr. Christopher Nelson for scientific editing. This work was supported by NIH/NINDS R01 NS100864 and start-up funds for DH from Departments of Oncological Sciences and Neurosurgery, Icahn School of Medicine, Mount Sinai. NIH/NINDS 1F31NS106887 provided funding for CH and NIH/NCI 1F31CA232531 provided funding for JR. DHG is supported by a Research Program Award from the National Institute of Neurological Disorders and Stroke (1R35NS097211).

## Author contributions

Conceptualization, D.H., Z.C.; Methodology: Z.C., D.H.; Investigation: Z.C., M.K., C.J.H., G.P., M.P.V., J.L.R., J.A., V.M., F.S., M.M., A. A., N.N.; Visualization, Z.C., B.G., S.C.; Formal Analysis, Z.C., B.G., M.K., S.C.; Resources, N.T. and D.S.; Software, Z.C., S.C., B. G., and A.T.; Writing-Original Draft, Z.C. and D.H.; Writing-Review & Editing, S.C., D.H.G., B.G., and A.T.; Funding Acquisition, D.H.; Supervision, D.H., A.T., and F.M.

## Declaration of interests

The authors have no relevant competing interests to disclose.

## Materials and Methods

### Mice

Mice of both sexes (equal distribution) in the age range of 8-16 weeks were used for experiments (Chen et al., 2020; Herting et al., 2017). Previously-described *Il1b^-/-^*, and *Il1a/b^-/-^* mice were crossed with the *Ntv-a* mice to generate *Il1b^-/-^;Ntv-a* and *Il1a^-/-^;Il1/b^-/-^;Ntv-a* mice (Glaccum et al., 1997; Herting et al., 2019; Horai et al., 1998). Mice used in this study are all in a C57BL/6 background, except for *Ntv-a;Cdkn2a^-/-^;Pten^fl/fl^*, which is in a mixed genetic background. C57BL/6J mice (000664) at 8-12 weeks old were purchased from the Jackson labs and used for BMDM isolation. All animals were housed in a climate-controlled, pathogen-free facility with access to food and water *ad libitum* under a 12-hour light/dark cycle. All experimental procedures were approved by the Institutional Animal Care and Use Committee (IACUC) of Emory University (Protocol #201700633) and the Icahn School of Medicine at Mount Sinai (Protocol #201900619).

### Virus generation and tumor induction

We delivered RCAS-*PDGFB* in *Cdkn2a^-/-^;Ntv-a* mice, or the combination RCAS-*PDGFB* with *RCAS-shRNA-p53-Rfp* in *Ntv-a* mice, both of which have been shown to closely resemble human PN GBM in terms of histological and molecular features (Chen et al., 2020; Herting et al., 2017). To propagate RCAS viral vectors, DF-1 cells (ATCC, CRL-12203) were purchased and grown at 39°C according to the supplier’s instructions. The cells were routinely tested for mycoplasma contamination to assure they are negative for mycoplasma. Cells were transfected with *RCAS-hPDGFB-HA* and RCAS-shRNA-*p53*-*Rfp* using a Fugene 6 transfection kit (Roche, 11814443001) according to the manufacturer’s instructions. DF-1 cells (4×10^4^) in 1 μl neurobasal medium were stereo-tactically delivered with a Hamilton syringe equipped with a 30-gauge needle for tumor generation (Franklin and Paxinos, 1997). The target coordinates were in the right-frontal striatum for PDGFB-overexpressing tumors at AP 1.0 mm and right 1.5 mm from bregma; depth 1.5 mm from the dural surface. *Nf1*-silenced tumors were generated via injection into the subventricular zone at coordinates AP 0.0 mm and right 1.0 mm from bregma; depth 1.5 mm from the dural surface (Chen et al., 2020; Franklin and Paxinos, 1997; Herting et al., 2017). Mice were continually monitored for signs of tumor burden and were sacrificed upon observation of endpoint symptoms including head tilt, lethargy, seizures, and excessive weight loss.

### Orthotopic glioma generation

The same procedure was used as described above, except 3×10^4^ of freshly-dissociated tumor cells were injected in the right-frontal striatum AP 1.0 mm and right 1.5 mm from bregma; depth 1.5 mm from the dural surface of recipient animals. Two or three donor tumors of either sex were used for obtaining single cell suspension for orthotopic glioma generation in male and female recipient animals.

### Monocyte-derived macrophage isolation and culture

C57BL6/J mice (8-12 weeks of age) were euthanized via CO_2_ asphyxiation. The whole, intact femur and tibia were stripped of muscle and collected in sterile Dulbecco’s phosphate-buffered saline (DPBS). Both ends of the bone were cut and the marrow was flushed into a clean petri dish using DPBS supplemented with 4% bovine serum albumin (BSA) (ThermoFisher, 15260037), heparin (StemCell, 07980), DNase I (Sigma-Aldrich, 11284932001), and penicillin/streptomycin (Fisher, SV30010). The resulting mixture was briefly triturated and passed through a 70 μm cell strainer. The cells were plated in a 15 cm non-cell culture-treated plate for a six-day differentiation in 15 ml of DMEM (ThermoFisher, 10569010) with 10% fetal bovine serum (FBS; HyClone, SH30396.03) and 40 ng/ml M-CSF (Peprotech, 315-02). An additional 15 ml of media with M-CSF was added after day three. The cells were harvested for experimentation via a 10-minute incubation in DPBS with 5mM EDTA on ice. Cells were plated in 6-well plates in 2 ml of culturing media with macrophage colony-stimulating factor (M-CSF) at a concentration of 20 ng/ml for experimentation.

### Organotypic tumor slice culture

*Ntv-a* mice harboring PDGFB-overexpressing and *p53-* silenced tumors were euthanized at endpoint via carbon dioxide asphyxiation. The brain was rapidly extracted and embedded in 4% low-melt agarose in PBS. The embedded brain was then mounted on a vibratome (Leica, 1220S) and submerged in ice cold Hank’s balanced salt solution (HBSS, Gibco 14185052). The brain was cut into 300 μm-thick sections and the slices were transferred to inserts in a 6-well plate. Slices were cultured in Neurobasal media (StemCell, 05700) supplemented with B27 supplement (ThermoFisher, 17504044), sodium pyruvate (ThermoFisher, 11360070), and glutamine (ThermoFisher, 35050061). M-CSF (BioLegend, 576406) at a concentration of 40 ng/ml was included in the media during coculture experiments with BMDM.

### Tumor and cultured cell RNA isolation and analysis

At endpoint, mice were sacrificed with an overdose of ketamine and immediately perfused with ice cold Ringer’s solution (Sigma-Aldrich, 96724-100TAB). The brain was extracted, and a piece of tumor was immediately snap-frozen in liquid nitrogen for storage at −80°C. Alternatively, cultured cells were harvested from plates using TRIzol (ThermoFisher, 15596026). RNA was isolated from the frozen tumor pieces or cells with the RNeasy Lipid Tissue Mini Kit (Qiagen, 74804) according to the manufacturer’s instructions. RNA quantity was assessed with a NanoDrop 2000 spectrometer (ThermoFisher), while quality was confirmed via electrophoresis of samples in a 1% bleach gel as previously described (Aranda et al., 2012). RNA was used to generate cDNA with a First Strand Superscript III cDNA synthesis kit (ThermoFisher, 18080051) according to the manufacturer’s instructions and with equal amounts of starting RNA. Quantitative-PCR was performed with the validated BioRad PCR primers using SsoAdvanced Universal green Supermix (BioRad, 1725271). Fold changes in gene expression were determined relative to a defined control group using the 2^-ΔΔCt^ method or by z-score, with β-Actin or HPRT used as housekeeping genes.

### Immunoblot analysis

Mouse primary GBM cell lines were treated with 100 pM IL-1β (R&D Systems, 401-ML/CF) for 10, 30, and 60 minutes. Cells were washed 2 x with ice-cold PBS and scraped into RIPA buffer (50 mM Tris, pH 8.0, 150 mM NaCl, 5 mM EDTA, 1% NP40 and 0.5% deoxycholate, 10 mM NaF, 1 mM sodium orthovanadate with complete protease inhibitor cocktail (Roche) as describe previously, incubated on ice for 30 min, centrifuged, and protein concentrations were determined using a BCA kit (Pierce, 23227). Cell lysates (10-50 μg) were subjected to SDS-polyacrylamide gel electrophoresis on a 4-20% precast gradient gels (BioRad, 5678094) and proteins were transferred to nitrocellulose membranes (Bio-Rad, 162-0112). After transfer, membranes were incubated in 5% Nonfat Dry Milk (Cell Signaling Technology, 9999) in 1 x TBST (Cell Signaling, 9997) for 1 hour. Membranes was incubated for 16 hours at 4°C with primary antibody:P-NF-kappa B p65 (1:1000), NF-kB (1:1000), P-IkB-alpha (1:1000), and IkB-alpha primary antibodies (1:1000); all antibodies were from the NF-kB Pathway Sampler kit (Cell Signaling, 9936S) and β-actin was from Abgent (1:5000, ABIN1842939) in TBST (Cell Signaling, 9997). Immunodetection was performed with anti-rabbit horseradish-peroxidase-conjugated secondary antibody (1:1000) and anti-mouse horseradish–conjugated secondary antibody (1:1000); secondary antibodies were from NF-kappa B Pathway Sampler kit (Cell Signaling, 9936S). Immunodetection was performed with Chemiluminescent HRP Antibody Detection Reagent (Denville Scientific INC, E2400).

### Human tissue samples and pathological appraisal

Archived formalin-fixed, paraffin-embedded (FFPE) human GBM samples and de-identified clinical information were provided by Emory University. Fresh tumor tissues used for ELISA quantification were collected at Mount Sinai Hospital through the biorepository, under IRB-approved protocols (18-00983). All patient samples were deidentified. Ten randomly selected tumor samples were included in this study, with de-identified age, sex, and molecular information summarized in Table S3. “Normal appearing” brain tissues away from the tumor mass, obtained during surgical resection, from three patient samples were used as controls. Board-certified neuropathologists diagnosed and graded both the human tumor tissues and murine samples according to the 2016 World Health Organization Classification of Tumors of the Central Nervous System (Louis et al., 2016). Gene expression profiling to determine transcriptional subtypes was performed using targeted genome sequencing (Foundation One/Sema4).

### TCGA analysis

U133 Microarray data for the GBM (TCGA, provisional) dataset were downloaded from cBioPortal (https://www.cbioportal.org) in August 2019 and sorted into subtypes based upon a proprietary key. G-CIMP-positive tumors were excluded from analysis. We included 372 patient samples for which covariate information (survival information, age, and gender) was available. Cox Proportional Hazard Models were fitted in R using age and gene expression as continuous covariates, and gender as a binary variable. Forest plots were done using the function *ggforest*.

### Tissue processing and immune-histochemistry

Archived FFPE human GBM samples were sectioned at 5 μm thickness, slide-mounted, and stored at −80°C until use. To process mouse tumor tissues, animals were anesthetized with an overdose of ketamine/xylazine mix and transcardially perfused with ice-cold Ringer’s solution. Brains were removed and processed according to the different applications. For H&E tumor validation and immunohistochemistry staining, brains were fixed in 10% neutral buffered formalin for 72 hours at room temperature (RT), processed in a tissue processor (Leica, TP1050), embedded in paraffin, sectioned (5 μm), and slide mounted.

Immunohistochemistry staining was performed on either Discovery XT platform (Ventana Medical Systems) or Leica Bond Rx (Leica). Primary antibodies used in this study include: anti-IBA-1 (1:1500, Wako, 019-19741), anti-phosphorylated Histone 3 (1:200, Millipore, 06-570), anti-CD44 (1:100, BD Pharmingen, 550538). Digital images of the slides were acquired by using a Nanozoomer 2.0HT whole-slide scanner (Hamamatsu Photonic K.K) and observed offline with NDP view2 software (Hamamatsu). Image analysis was performed using Fiji (NIH).

### Tumor dissociation and primary cell culturing

Tumor dissociation was performed as previously described. Briefly, tumors were dissected from the brain, minced into pieces < 1 mm^3^, and digested with an enzymatic mixture that includes papain (0.94 mg/ml, Worthington, LS003120), EDTA (0.18 mg/ml, Sigma, E6758), cysteine (0.18 mg/ml, Sigma, A8199), and DNase (60 μg/ml, Roche, 11284932001) in 2 ml HBSS (Gibco, 14175-095). Tumor tissues were kept at 37°C for 30 minutes with occasional agitation. The digestion was terminated with the addition of 2 ml Ovomucoid (0.7 mg/ml, Worthington, LS003086). Following digestion, single cells were pelleted, resuspended in HBSS, and centrifuged at low speed (84 RCF) for 5 min, before passing through a 70 μm cell strainer.

For tumorsphere cultures (NSC), cells were seeded at 5 × 10^5^ cells/ml and grown in Neurocult mouse neurobasal medium (Stem Cell Technologies, 5700) supplemented with 10 ng/ml hEGF (Lonza, cc-4017FF), 20 ng/ml basic-hFGF (ThermoFisher, PHG0261), 1 mg/ml Heparin (Stem Cell Technologies, 7980), and NSC Proliferation Supplements (Stem Cell Technologies, 5701). Fresh medium was added to the cultures every 48 hours. For differentiation (FBS) cultures, cells were seeded at 5 × 10^5^ cells/ml and grown in DMEM (ATCC, 30-2002) supplemented with 10% FBS (ATCC, 30-2020). Fresh medium was added to the cultures every 48 hours.

### IL-1β treatment *in vitro*

For primary neurospheres, cultures (before rIL-1β stimulation) were dissociated with Accutase (Sigma-Aldrich, A6964) to generate single cells, which were then grown as adherent monolayers on coverslips (Φ=12 mm) coated with Geltrex (Life Technologies, A14132-01) prepared according to manufacturer’s instructions. Cells were stimulated with 100pM rIL-1β (R&D 401-ML/CF) for period indicated on the graphs.

### Immunofluorescence

Human GBM 5 μm FFPE sections were stained with anti-IBA1 (1:500, Wako, 019-19741) and anti-IL-1β (NCI preclinical repository, biological resource branch, 32D). Secondary antibodies conjugated to Alexa-Fluor dyes (555 nm, 647 nm from Invitrogen) at a dilution of 1:500 in PBS/2% BSA were applied. DAPI (Sigma, D9542) was used for nuclear counterstaining. Autofluorescence from erythrocytes was recorded in the green emission channel. Fluorescence images were taken on an Olympus FV1000 confocal microscope and analyzed with FIJI software (NIH).

### Enzyme-linked immunosorbent assay

Cell lysates for enzyme-linked immunosorbent assay (ELISA) were collected via sonication of cells in lysis buffer supplemented with protease and phosphatase inhibitors. Tissue lysates were collected via mechanical homogenization in lysis buffer followed by sonication. Protein concentrations were determined using a Bradford protein assay (Bio-Rad, 5000001) according to the manufacturer’s instructions. ELISAs were performed for hIL-1β (R&D, DY201-05), mIL-1β (R&D, DY401-05), CCL2 (R&D, DY479), CCL7 (Boster Bio, EK0683), CCL8 (R&D, DY790), CCL12 (R&D, MCC120), and VEGF (R&D, MMV00) on cell lysates and cell supernatants according to the manufacturer’s instructions.

### Flow Cytometry and spectral flow cytometry

Initial steps of the enzymatic dissociation of the tumors are the same as described above, except 0.5% collagenase D (Sigma, 11088858001) and DNase (Roche, 11284932001) were used in place of papain. Single-cell suspensions were passed through 70μm cell strainers, centrifuged, and resuspended in 30% Percoll (GE Healthcare, 17-0891-01) solution containing 10% FBS (Hyclone SH30396.03). Cells were separated by centrifugation at 800*g* for 15 minutes at 4°C. The supernatant was carefully removed to discard debris and lipids. The cells were then washed in cold PBS and resuspended in RBC lysis buffer (BioLegend, 420301) for 1 min at 37°C. Cells were transferred to an Eppendorf tube and washed once with FACS buffer (DPBS with 0.5% BSA) and blocked with 100 μl of 2x blocking solution (2% FBS, 5% normal rat serum, 5% normal mouse serum, 5% normal rabbit serum, 10 μg/ml anti-FcR (BioLegend, 101319) and 0.2% NaN3 in DPBS) on ice for 30 minutes. Cells were then stained on ice for 30 minutes and washed with PBS. The cells were subsequently incubated in 100 μl viability dye (Zombie UV, BioLegend, 1:800) at room temperature for 20 min. The cells were washed and fixed with fixation buffer (eBioscience, 005123-43, 00-5223-56) for 30 min. Cells stained with the cocktail of antibodies examined myeloid lineage are set aside in the fridge until loading to the cytometer. Cells stained for the lymphoid panel were then permeabilized with a permeabilization buffer (eBioscience, 00-8333-56) before the intracellular markers were stained. The cells were washed and stored in fridge till analysis. Antibodies used in this study include are listed in table S2. All data were collected on a BD LSR II flow cytometer or Cytek Aurora spectral flow cytometer. Data were analyzed off line using FlowJo 10 software (Tree Star Inc.).

### Single-cell RNA-seq and data analysis

Single cell suspensions of the tumors were obtained by papain dissociation as described above. Viability of single cells was assessed using Trypan Blue staining, and debris-free suspensions of >80% viability were deemed suitable for single cell RNA Seq. Samples with lower viability were run with caution. Single cell RNA Seq was performed on these samples using the Chromium platform (10X Genomics) with the 3’ gene expression (3’ GEX) V3 kit, using an input of ~10,000 cells. Briefly, Gel-Bead in Emulsions (GEMs) were generated on the sample chip in the Chromium controller. Barcoded cDNA was extracted from the GEMs by Post-GEM RT-cleanup and amplified for 12 cycles. Amplified cDNA was fragmented and subjected to end-repair, poly A-tailing, adapter ligation, and 10X-specific sample indexing following the manufacturer’s protocol. Libraries were quantified using Bioanalyzer (Agilent) and QuBit (ThermoFisher) analyses and were sequenced in paired end mode on a NovaSeq instrument (Illumina) targeting a depth of 50,000-100,000 reads per cell.

Raw fastq files were aligned to mouse genome reference mm10 customized to include the Rfp sequence, using CellRanger v5.0.0 (10X Genomics). Count matrices filtered by CellRanges algorithm were further filtered by discarding cells with either < 400 genes, < 1000 UMI, or > 25% mitochondrial genes expressed. Data was processed and analyzed using R package Seurat v4.0.5. Normalization was performed using NormalizeData function with normalization.method = ‘LogNormalize’. Dimensionality reduction was computed on the top 2,000 variable features using FindVariableFeatures, ScaleData and RunPCA functions. UMAPs were generated using the top 15 PCs. For subclustering the immune compartment, we used R package Harmony to mitigate for batch effects driven by technical variation between replicates. *De novo* clustering using the Louvain algorithm was applied at different resolutions (0.2; 0.8; 2; 5) on the KNN graph space. For high-level annotation, cell types were identified in an iterative and semi-supervised fashion by assigning de novo discovered clusters to cell types based on expression of known marker genes that define each cluster. Annotation of cell subtypes at a lower-level was performed in a similar manner as for the high-level and further aided by *de novo* marker discovery using the Seurat FindMarkers function and Wilcoxon Rank Sum test. To identify doublet-enriched clusters we looked for clusters of cells displaying expression of canonical markers for two or more different cell types and higher number of genes/UMI; such clusters were removed from further analysis. Cell-level proliferation analysis was carried out with Seurat function CellCycleScoring using a list of murine cell cycle genes. Cells were then assigned as ‘cycling’ or ‘non-cycling’ if either their S-phase score or G2M-score was above a threshold of 0.05 for the tumor compartment and 0.1 for the immune compartment.

Identification of modules of co-expressed genes was carried out using the R package scWGCNA (https://github.com/smorabit/scWGCNA) by first computing metacells of 100 cells (k=100) using function construct_ metacells. To identify modules, function blockwiseConsensusModules was called with following parameters: softPower=12, deepSplit=3, mergeCutHeight = 0.25. Only the top 2,000 variable genes were used.

### Hematoxylin and eosin tumor volume reconstruction

Mice were sacrificed 25 days post-tumor cell transplantation with an overdose of ketamine and xylazine and perfused with 4% paraformaldehyde in PBS. The brain was carefully extracted and incubated in 4% paraformaldehyde in PBS for 24 hours following a 72-hour incubation in 30% sucrose in PBS. The brain was then embedded in O.C.T. compound (VWR, 25608-930) and frozen on dry ice. The entire brain was then serially sectioned on a cryostat (Leica) set to cut 30 μm sections. Every tenth section was collected and mounted on a slide for automated hematoxylin and eosin staining as described above. The slides were scanned at 20x magnification with a whole-slide scanner (Hamamatsu). Tumor area in each section was determined in a blinded fashion in NDP.view2 and multiplied by the thickness of ten slices. The resulting volumes for the slides of each tumor were then summed, producing an estimation of the total volume.

### Cannula installation and drug administration in tumor-bearing mice

Mice were anaesthetized with a ketamine (100 mg/kg) and xylazine (10 mg/kg) cocktail, their heads shaved, and they were placed into a stereotactic device with a three-axis micromanipulator. A burr hole was made with a compact drill above the hemisphere ipsilateral to the tumor at 1 mm lateral, −0.5 mm caudal relative to the bregma. A guide cannula (Plastics One) was implanted into the brain through the craniotomy and fixed in place with dental acrylic. A matching stylet or “dummy-cannula” was then screwed onto the guide cannula to prevent back flowing of the CSF or environmental debris from entering the guide cannula. Once the dental acrylic is cured, the skin around the pedestal of the guide cannula was sealed with Gluture liquid stitches (World Precision Instruments, 503763).

On the day of the infusion, the mice were anesthetized with inhalable isoflurane at 3% (volume/volume in pure oxygen) with a nebulizer. The dummy cannula was removed, and a Hamilton syringe mounted on a stereotactic frame was lowered into the guide cannula. Up to 2 μl of vehicle (artificial CSF, Harvard Instrument), IL-1RA (US Biological Corporation, I7663-62E), or purified anti-IL-1β IgG (BioXcell, BE0246) were injected into the lateral ventricle over a 1 min. Once the injection was completed, the dummy cannula was screwed back, and mice were returned to their home cages.

### Statistical analyses

Graphs were created using GraphPad Prism 8 (GraphPad Software Inc.) or R. Variables from two experimental groups were analyzed using unpaired or paired parametric Student’s two-tailed *t*-tests as appropriate, assuming equal standard deviations. One-way ANOVA was used to compare variables from more than two groups. Kaplan–Meier survival analysis was used to analyze survival difference in mGBM using the log-rank (Mantel-Cox) test and Gehan-Breslow-Wilcoxon test. Data are shown as mean±S.D. Number of samples in each group is indicted in the individual figures. (*) *P* < 0.05; (**) *P* < 0.01; (***) *P* < 0.001; (****) *P* < 0.0001; (ns) not significant.

## Data availability statement

The data that support the findings of this study are available from the corresponding author upon reasonable request. ScRNA-Seq data were deposited at TBD. Code used to produce analyses and figures is available at GitHub (TBD).

## Notes

### Competing Interest Statement

The authors have declared no competing interest.

