## Supplemental figures for "A paracrine circuit of IL-1β/IL-1R1 between myeloid and tumor cells drives glioblastoma progression"

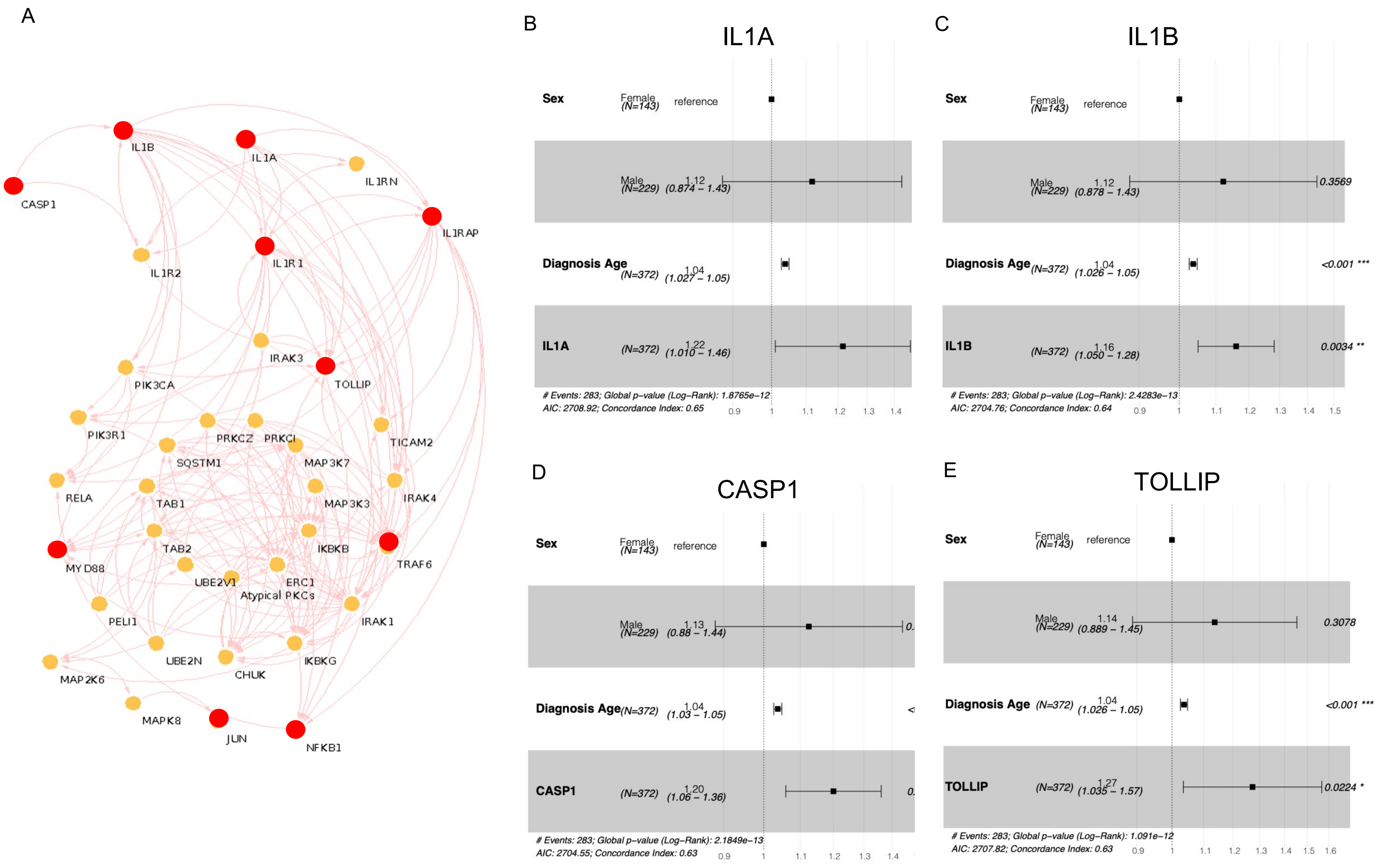


**Supplementary Figure 1. Association between the expression of *IL-1* pathway and the survival time of IDH-WT GBM patients**. **(A)** Molecules of the *IL-1* signaling pathway. Red dots highlight selected molecules that are prominent players in this pathway. (**B**) – **(E)** Forrest plots generated using Cox Proportional Hazards models, using expression of different genes (indicated) as continuous covariates.

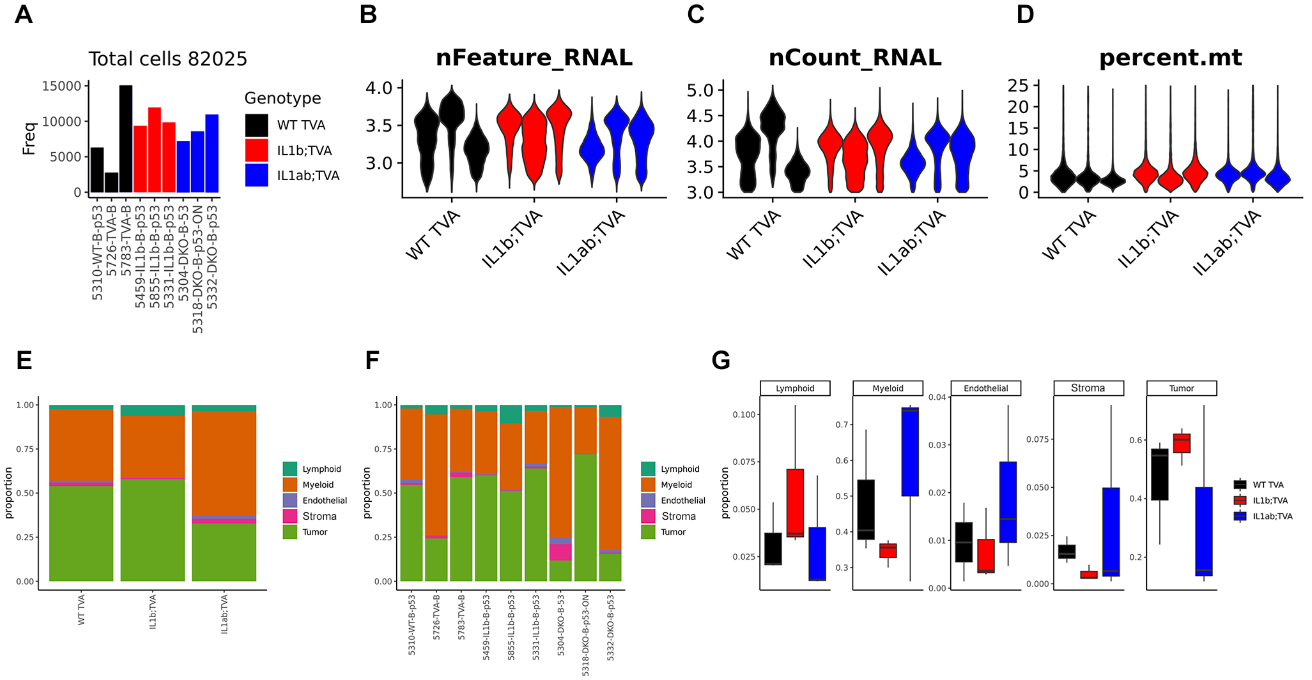


**Supplementary Figure 2. Single cell RNA seq analysis of tumors generated in *WT;Ntv-a*, *Il1b$^{-/-}$;Ntv-a* and *Il1a$^{-/-}$;Il1b$^{-/-}$;Ntv-a* mice.** (**A**) Total number of cells per samples after removing doublets. (**B**) Distribution of number of unique molecular identifier (UMI) per cells per sample. (**C**) Distribution of number of genes detected per cell per sample. (**D**) Distribution of percentage of mitochondrial genes per cell per sample. (**F**) Proportion of the five major cell classes grouped individual samples. (**G**) boxplot showing the distribution of the five major cell classes in the three genotypes.

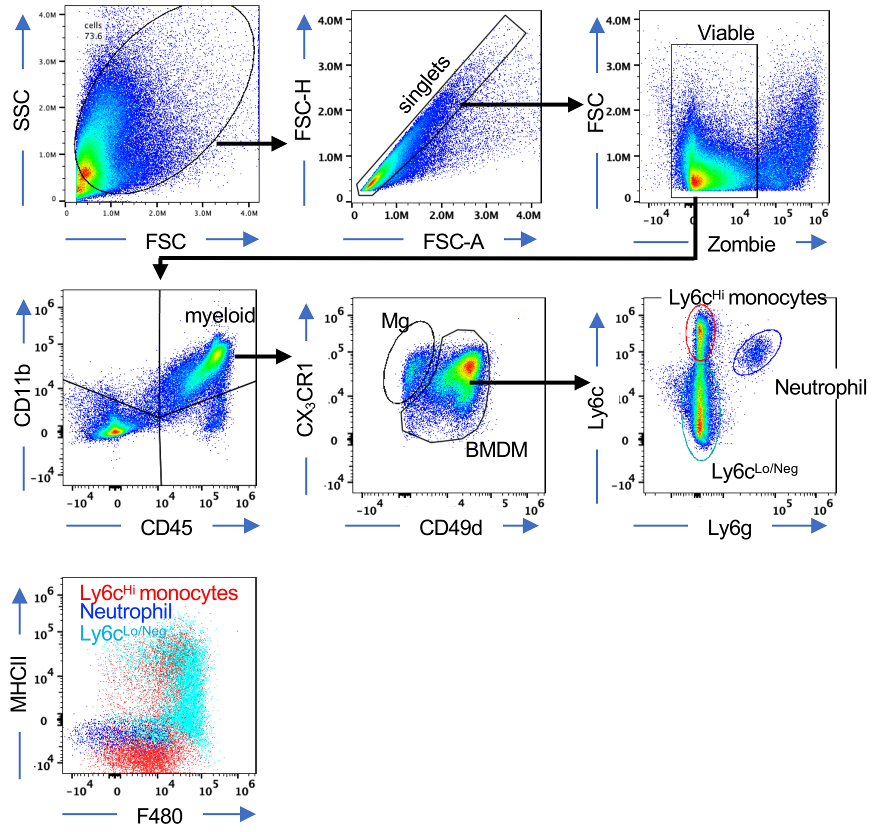


**Supplementary Figure 3. Gating strategy for multiplex Aurora spectral flow cytometry panel for myeloid cell subsets.** Mg= microglia, BMDM= bone marrow derived myeloid cells.

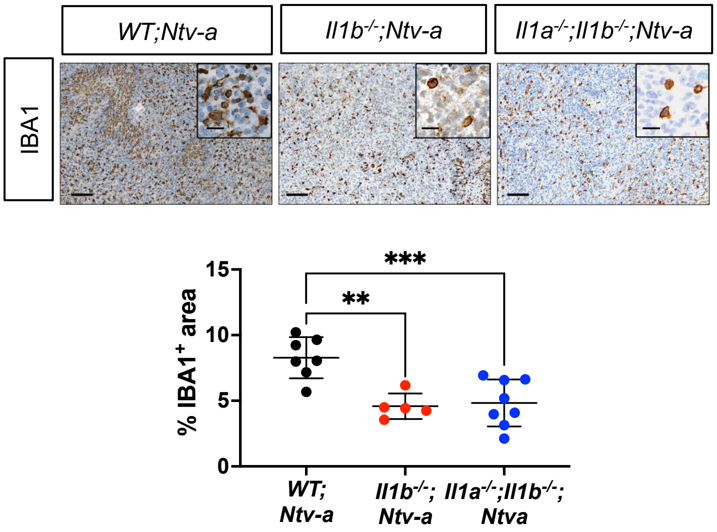


**Supplementary Figure 4. IHC quantification for CD44.** Scale bar = 50 μm; scale bar of inset = 20 μm. One way ANOVA with Tukey's *post-hoc* test., **P<0.01, ***P<0.001.

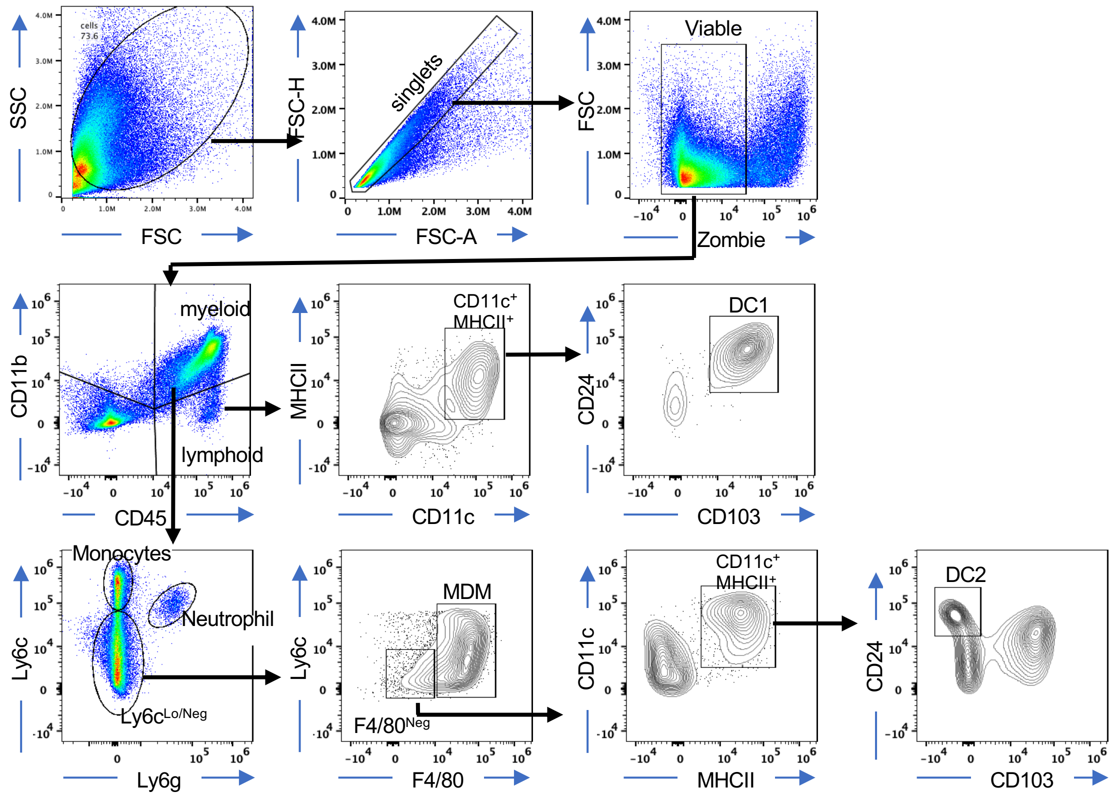


**Supplementary Figure 5. Gating strategy for multiplex Aurora spectral flow cytometry panel of DC1 and DC2.** MDM: monocyte-derived macrophage.

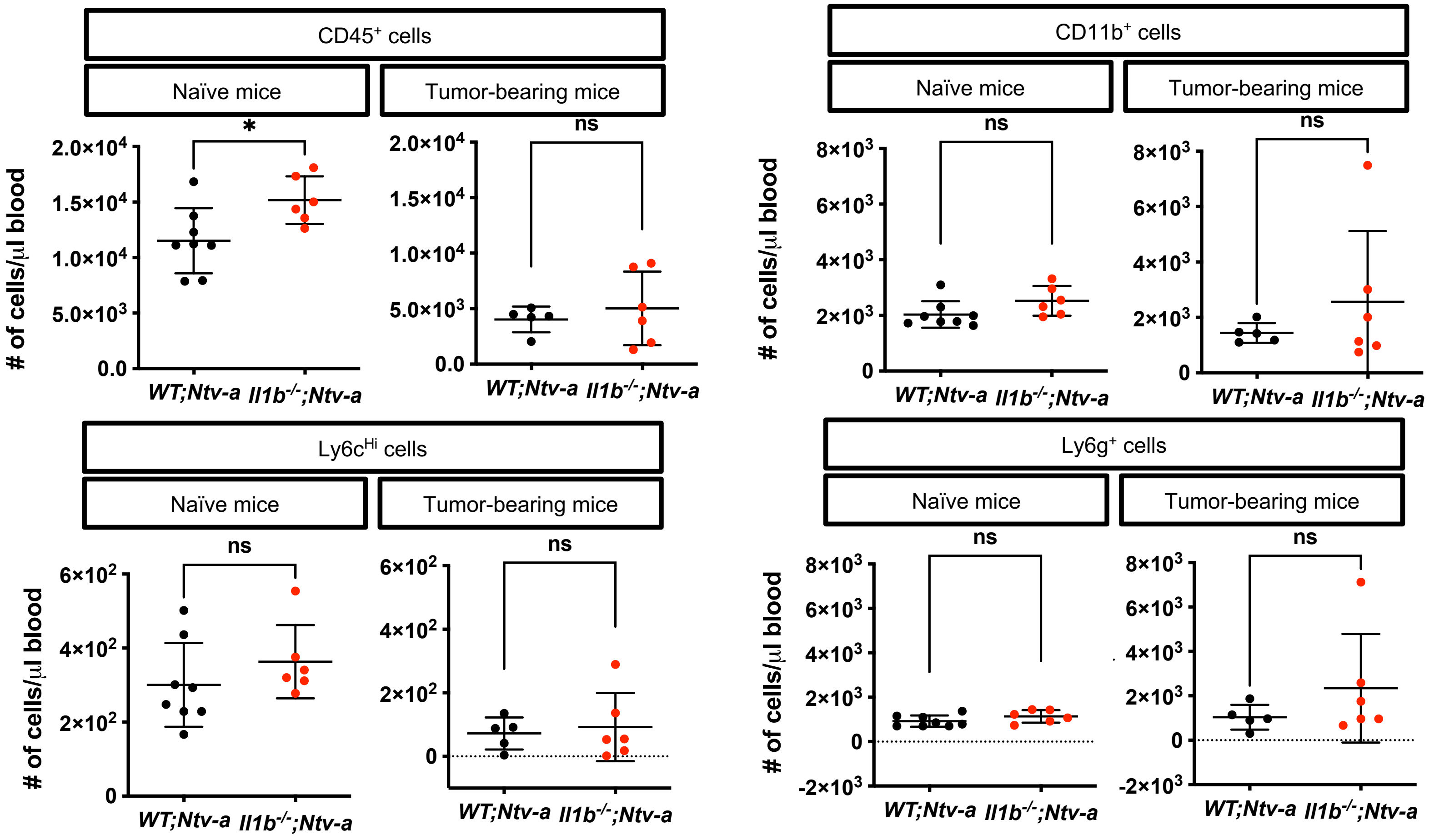


**Supplementary Figure 6. *Il1b* ablation has no impact on myeloid composition in the blood of naïve or tumor-bearing mice.** Quantification of FACS analysis of various myeloid cell populations in naïve and *PDGFB*-driven tumor-bearing mice in *WT;Ntv-a* and *Il1b$^{-/-}$;Ntv-a* mice. Two-tailed Student's *t*-test, *P<0.05, ns=not significant.

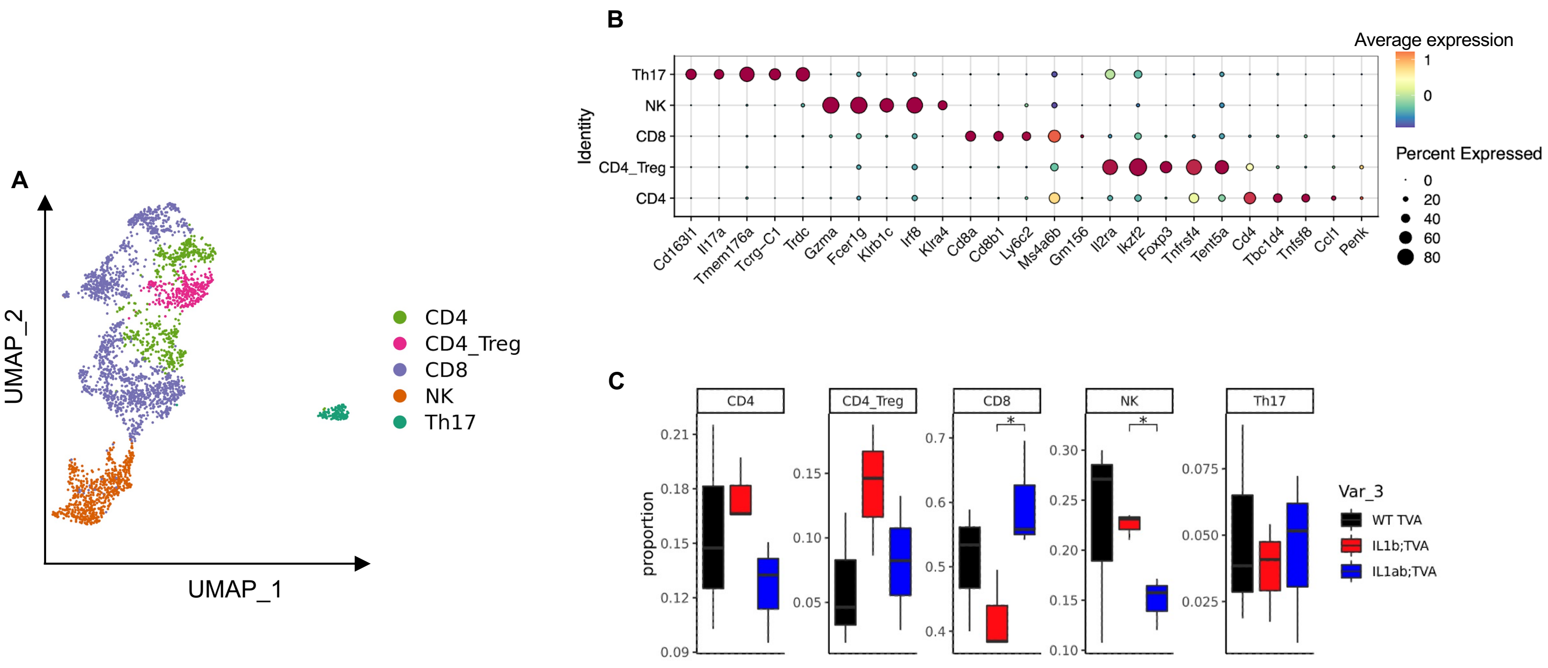


**Supplementary Figure 7. ScRNA-seq analysis for the lymphoid cells. (A)** UMAP dimensionality reduction of the lymphoid cell. (**B**) Selected marker genes used to annotate the cell types. (**C**) Lymphoid cell subtype distribution in *WT;Ntv-a*, *$Il1b^{-/-}$;Ntv-a* or *$Il1a^{-/-}$;$Il1b^{-/-}$;Ntv-a* mice.

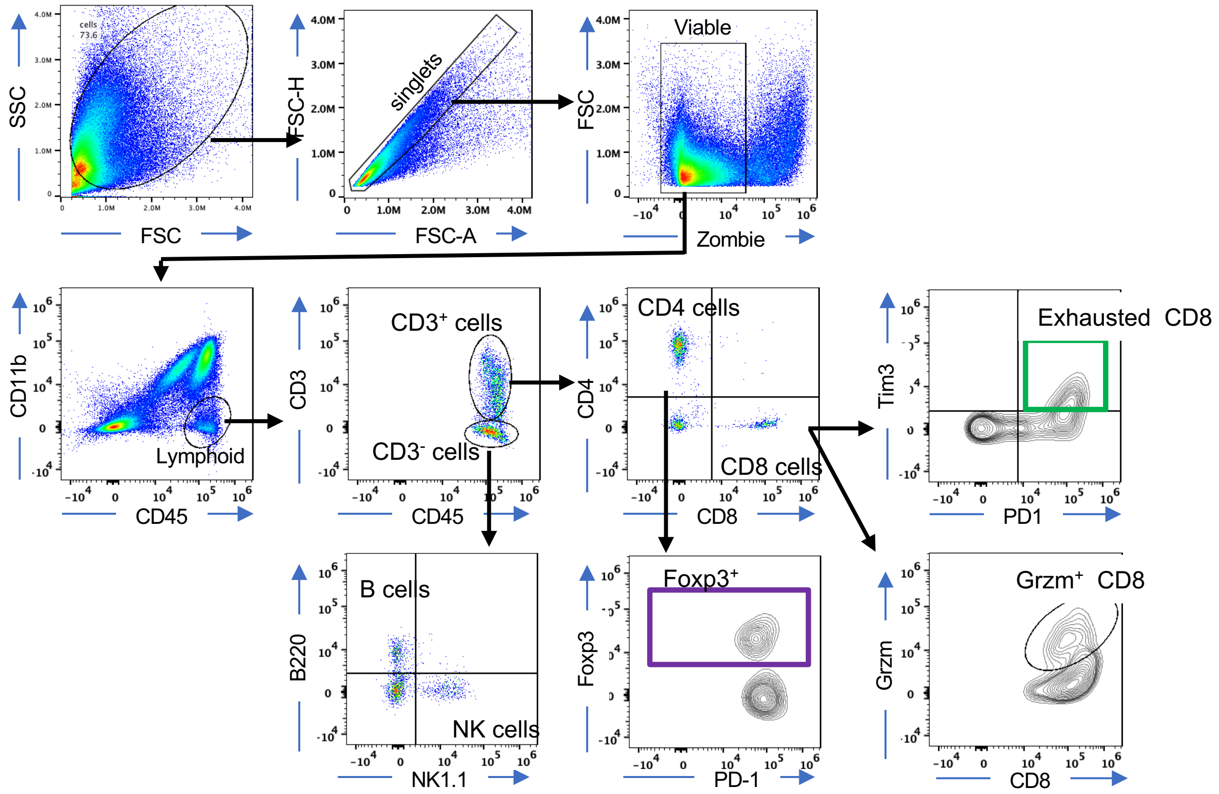


**Supplementary Figure 8. Gating strategy for multiplex Aurora spectral flow cytometry panel for lymphoid subsets.** Grzm=granzyme B.

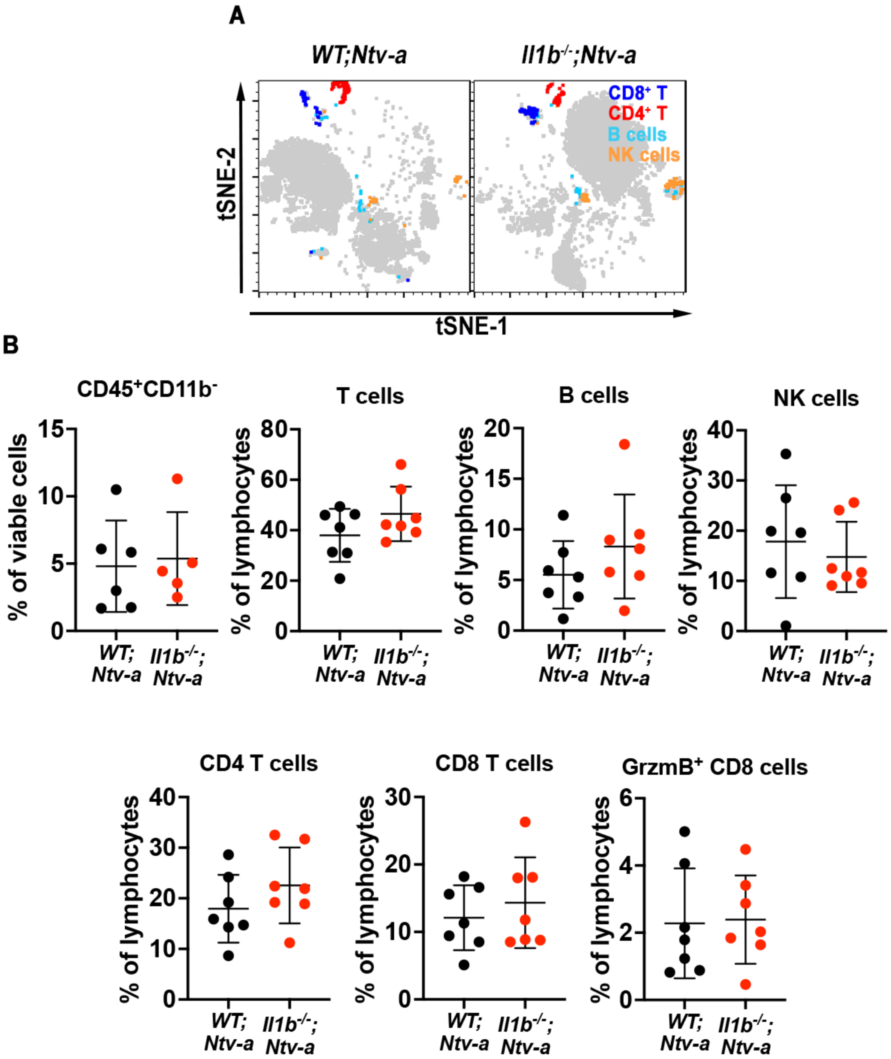


**Supplementary Figure S9.** ***Il1b*** **ablation has no impact on the total number of lymphoid cells in** ***PDGFB*****-driven murine GBM. (A)** tSNE plots illustrating the tumor cell/lymphoid composition in *WT;Ntv-a* and *Il1b$^{-/-}$;Ntv-a* mice bearing *PDGFB*-driven GBM. (**B**) Quantification of various lymphoid populations in tumors generated in *WT;Ntv-a* and *Il1b$^{-/-}$;Ntv-a* mice.

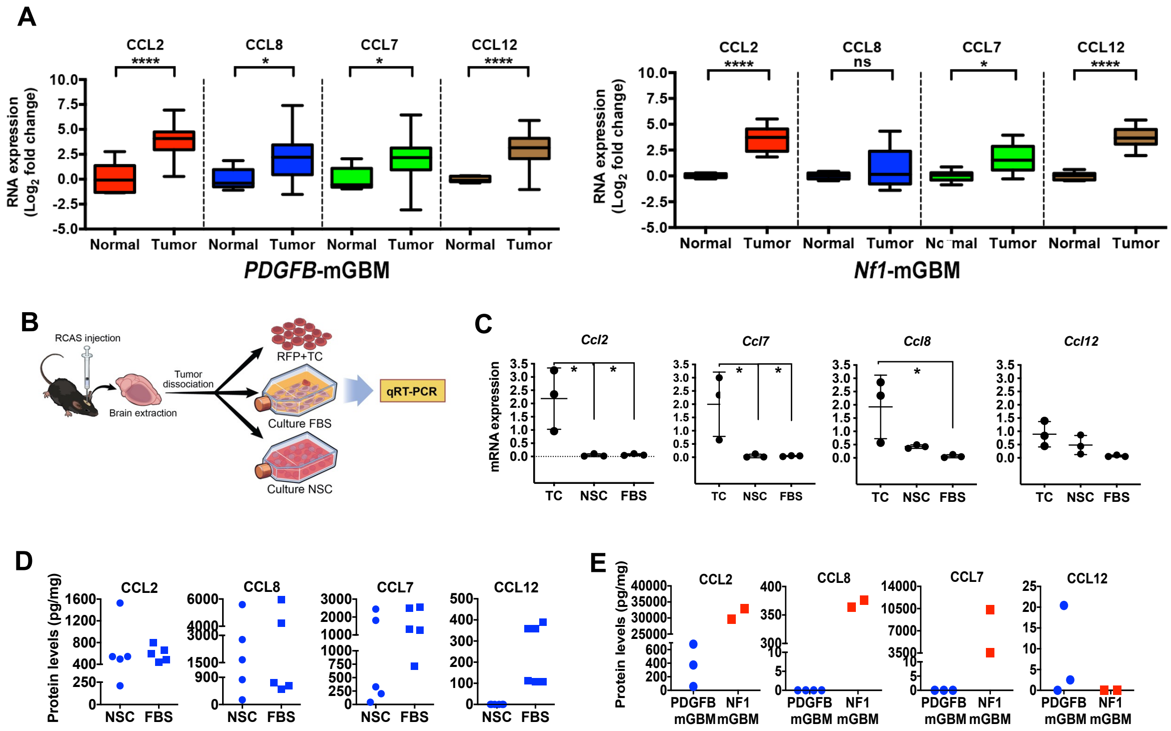


**Supplementary Figure S10. MCP expression is increased in both *PDGFB*-driven and *Nf1*-silenced tumors, and is induced by IL-1β stimulation *in vitro*. (A)** qPCR analysis of MCP family members - *Ccl2, Ccl8, Ccl7* and *Ccl12* - in naïve brain (normal), *PDGFB*-driven and *Nf1*-silenced murine GBM tumors. Two-tailed Student's *t*-test, *P<0.05, ****P<0.0001, ns=not significant. **(B)** Schematic illustration of experimental steps for **(C)** demonstrating that freshly-sorted tumor cells express higher levels of MCP proteins. When maintained in FBS-containing or NSC medium, expression levels decrease. One-way ANOVA, Dunnett's multiple comparison test, *P<0.05. **(D)** Expression of MCP proteins in early passage (P1-P2) of *PDGFB*-driven GBM cells grown in NSC or FBS conditions and **(E)** in late passage (P5 and above) *PDGFB*-driven GBM and *Nf1*-silenced GBM cells under FBS conditions. Each dot represents an independent primary culture.

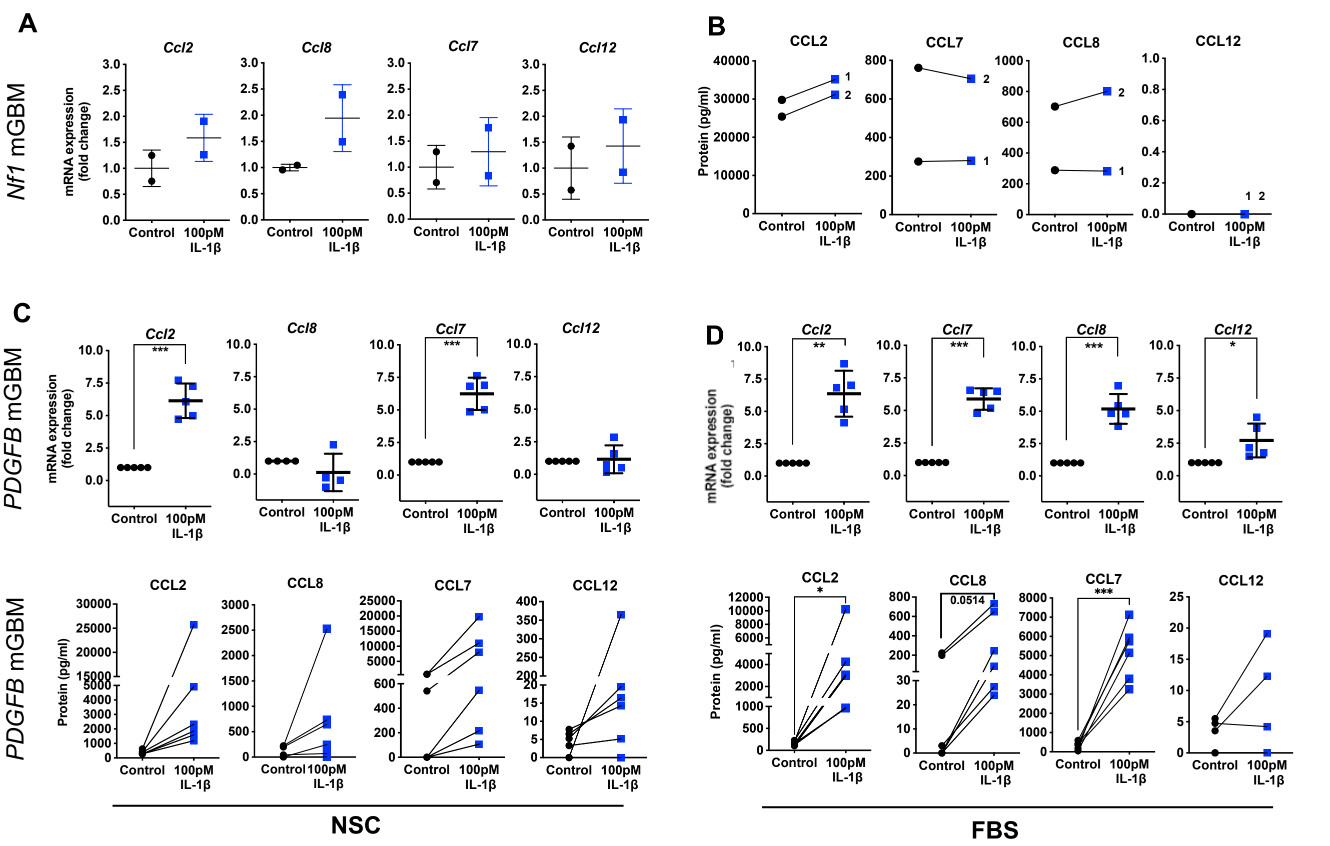


**Supplementary Figure S11. IL-1β stimulation induces the MCP network in *PDGFB*-driven, but not *Nf1*-silenced, mouse GBM cultures *in vitro*.** **(A)** IL-1β stimulation does not induce *Mcp* RNA or **(B)** protein expression in *Nf1*-silenced GBM cultures. Under **(C)** NSC or **(D)** FBS growth conditions, *PDGFB*-driven primary GBM cells respond to rIL-1β stimulation by increasing MCP network RNA and protein expression, as measured by qPCR and ELISA. Two-tailed Student's *t*-test, *P<0.05, **P<0.01, ***P<0.001.

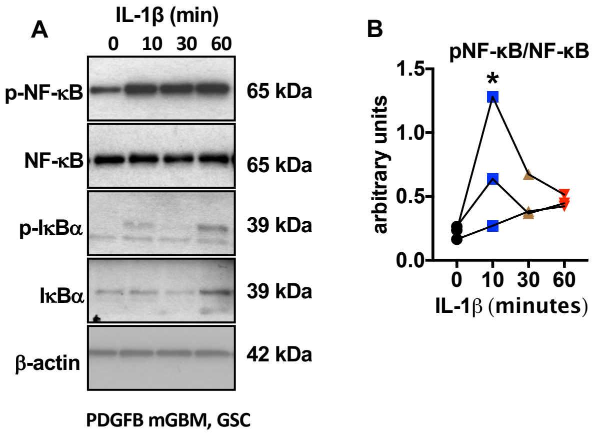


**Supplementary Figure 12. IL-1β stimulation induces NFκB pathway activation in *PDGFB*-driven, but not *Nf1*-silenced, mouse GBM cultures**. **(A)** Representative immunoblot showing NFκB pathway activation in *PDGFB*-driven mGBM cultures under NSC conditions following rIL-1β treatment *in vitro.* (**B**) quantification of the blots (n = 3). *P<0.05 by ANOVA test.

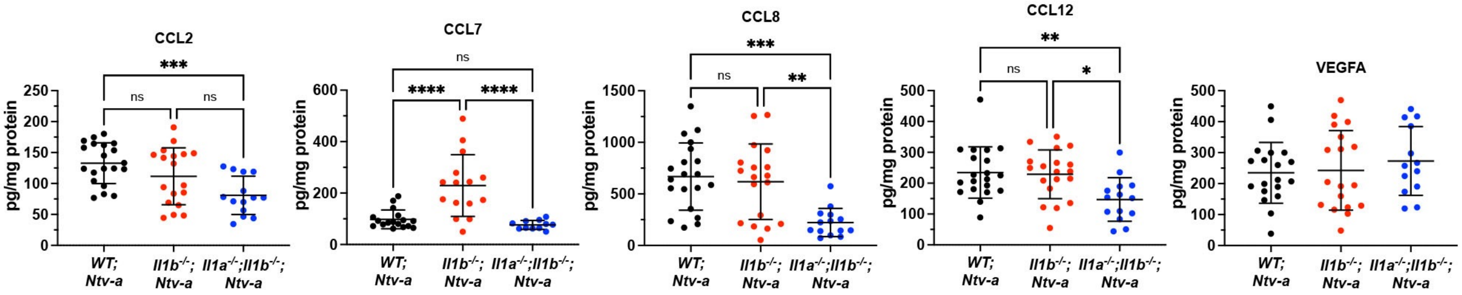


**Supplementary Figure S13. *Il1* genetic deletion reduces MCP production in *PDGFB*-driven GBM *in vivo***. Intracellular levels of MCP members (CCL-2, -7, -8, and -12) and VEGFA were quantified by ELISA in tumors generated in *WT;Ntv-a*, *Il1b$^{-/-}$;Ntv-a* or *Il1a$^{-/-}$;Il1b$^{-/-}$;Ntv-a* mice at the endpoint. VEGFA levels were not affected and were used as a negative control. One-way ANOVA with Tukey's post-hoc comparisons. *P<0.05; **P<0.01; ***P<0.001; ****P<0.0001; ns=not significant.

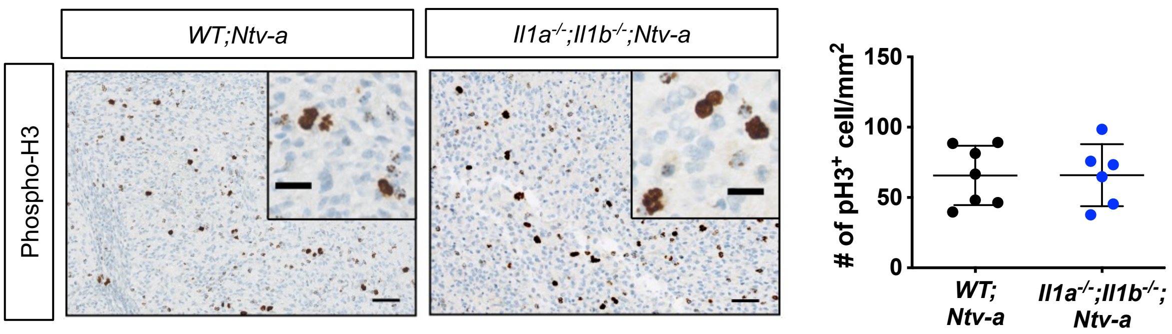


**Supplementary Figure 14. Immunohistological quantification of phosphorylated histone H3 in *WT;Ntv-a* and *Il1a*$^{-/-}$*; Il1b*$^{-/-}$*;Ntv-a* mice.** Scale bar = 50 μm; scale bar of inset = 20 μm. Student's *t*-test was used for statistical comparison.

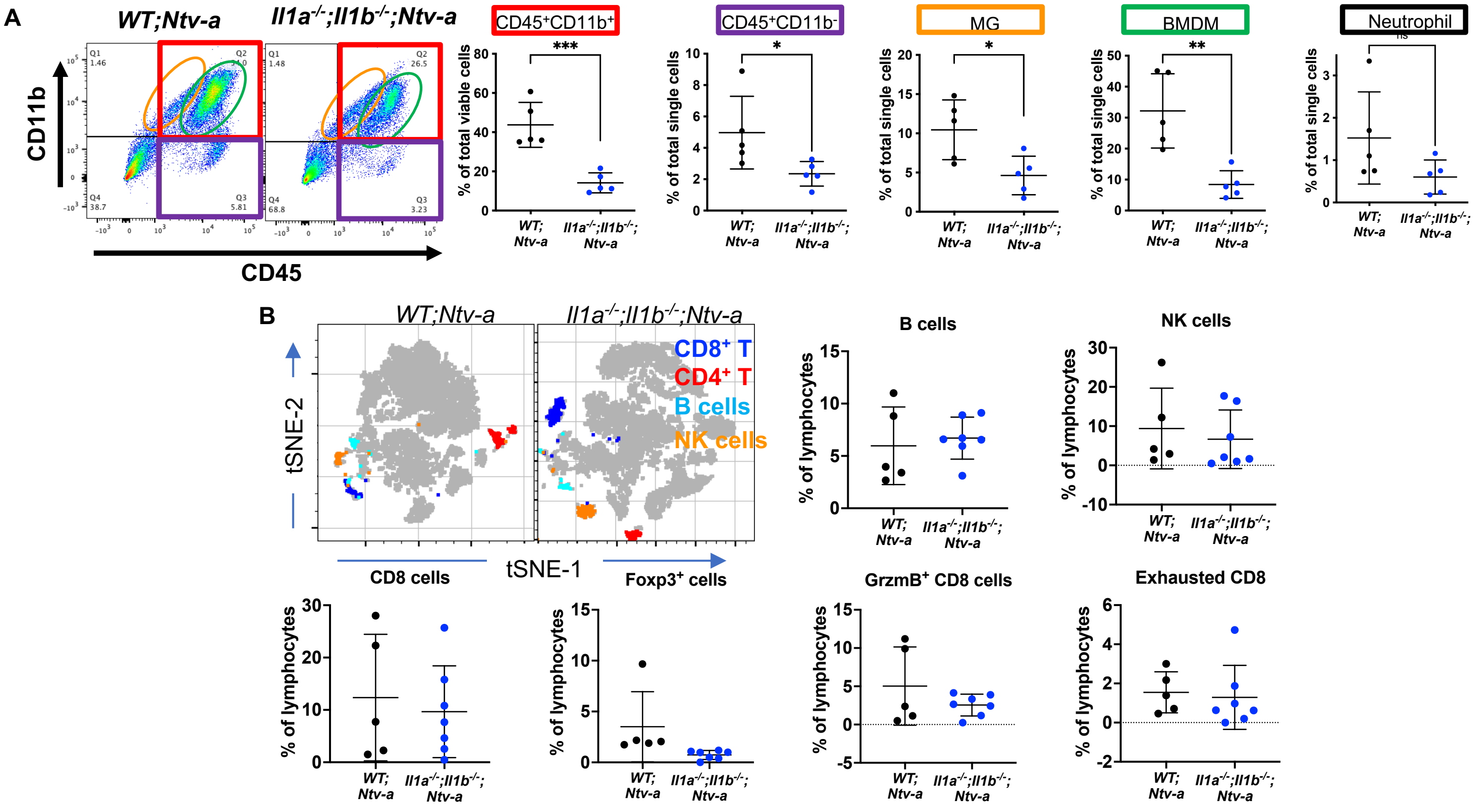


**Supplementary Figure 15. Flow cytometry and scRNA-seq analyses of lymphocytes in *WT;Ntv-a* and *$Il1b^{-/-}$;Ntv-a* mice. (A)** Flow cytometry dot plots and quantification of lymphocytes between *WT;Ntv-a* and *$Il1b^{-/-}$;Ntv-a* mice. Two-tailed Student's *t*-test, *P<0.05, **P<0.01; ***P<0.001. (**B**) tSNE plots and quantification of lymphocytes between *WT;Ntv-a* and *$Il1b^{-/-}$;Ntv-a* mice. Two-tailed Student's *t*-test.

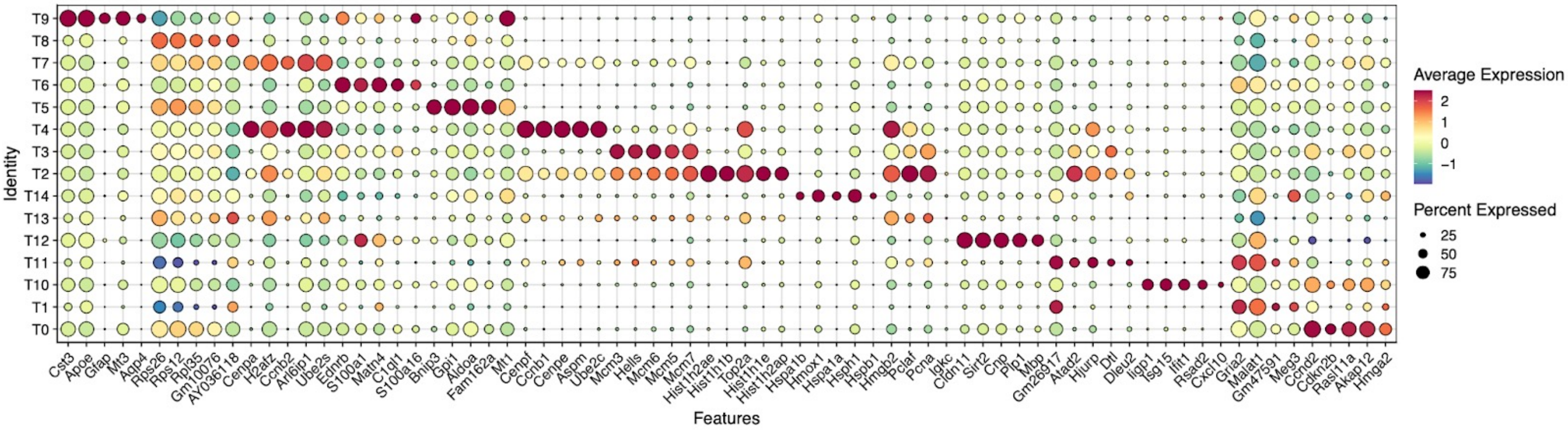


**Supplementary Figure 16. Signature gene sets used to cluster the malignant tumor cells.**

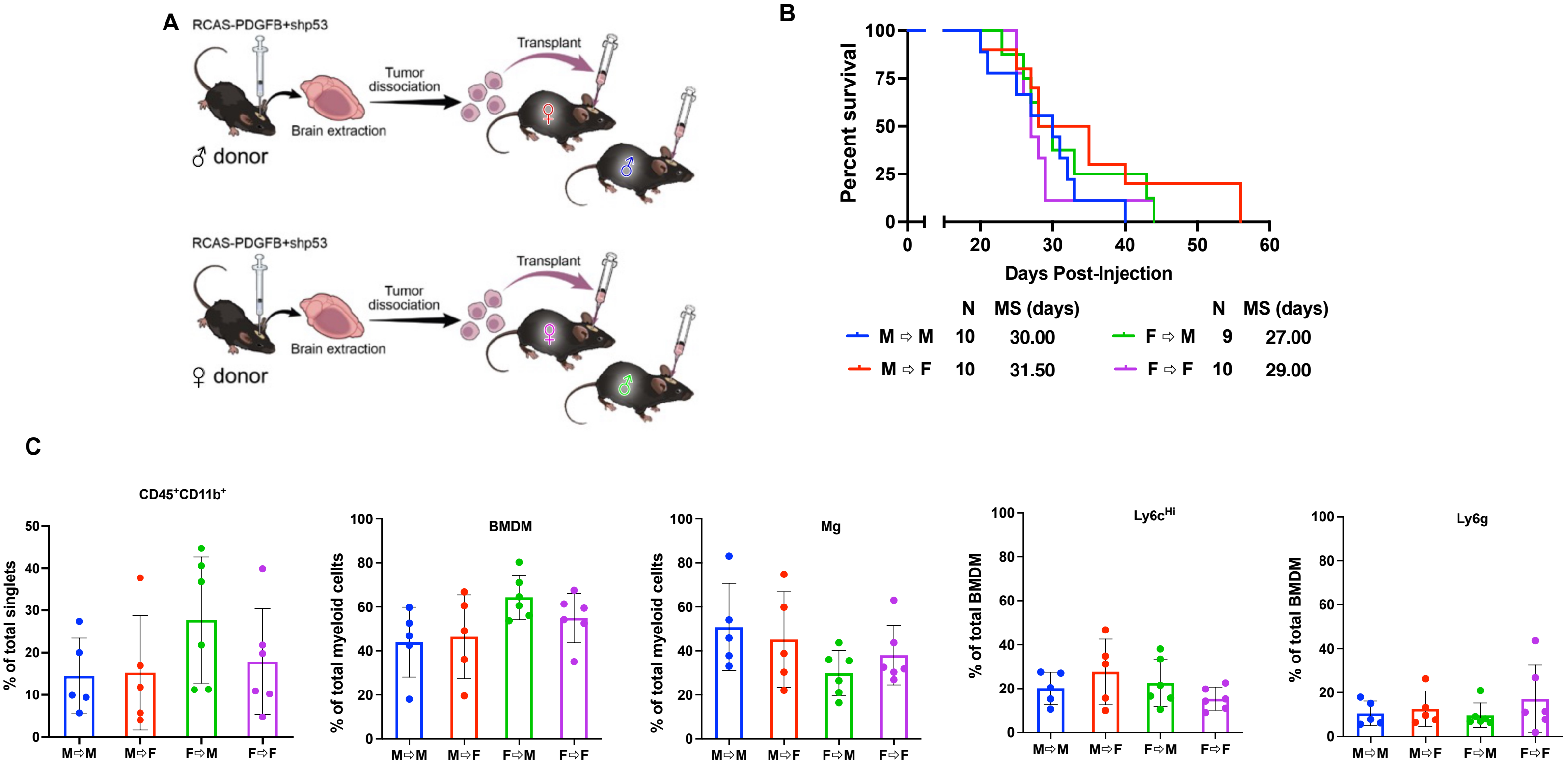


**Supplementary Figure 17. The sex of the tumor donor or the recipient animal was not associated with differential recruitment of myeloid populations and did not impact survival in PDGFB-tumor-bearing mice. (A)** Schematic illustration of the generation of various combinations of different sexes of donor and recipient mouse strains by orthotopic transplantation. **(B)** Kaplan-Meier survival curves. **(C)** Quantification of FACS analysis of various myeloid cell populations in female and male recipient animals following transplantation with male and female donor-derived tumor cells.
